# Hallmarks of slow translation initiation revealed in mitochondrially localizing mRNA sequences

**DOI:** 10.1101/614255

**Authors:** Thomas M. Poulsen, Kenichiro Imai, Martin C. Frith, Paul Horton

## Abstract

The mRNA of some, but not all, nuclear encoded mitochondrial proteins localize to the periphery of mitochondria. Previous studies have shown that both the nascent polypeptide chain and an mRNA binding protein play a role in this phenomenon, and have noted a positive correlation between mRNA length and mitochondrial localization. Here, we report the first investigation into the relationship between mRNA translation initiation rate and mRNA mitochondrial localization. Our results indicate that translation initiation promoting factors such as Kozak sequences are associated with cytosolic localization, while inhibiting factors such as 5′ UTR secondary structure correlate with mitochondrial localization. Moreover, the frequencies of nucleotides in various positions of the 5′ UTR show higher correlation with localization than the 3′ UTR. These results indicate that mitochondrial localization is associated with slow translation initiation. Interestingly this may help explain why short mRNAs, which are thought to initiate translation rapidly, seldom localize to mitochondria. We propose a model in which translating mRNA has reduced mobility and tends not to reach mitochondria. Finally, we explore this model with a simulation of mRNA diffusion using previously estimated translation initiation probabilities, confirming that our model can produce localization values similar to those measured in experimental studies.

## INTRODUCTION

Mitochondria most likely arose when oxygen-metabolizing bacteria were engulfed by a predatory ancestral eukaryotic cell and established a symbiotic relationship in which the bacteria supplied energy in return for shelter (1). Mitochondria subsequently lost the vast majority of their genomes (2) and therefore require imported proteins for their biogenesis and function. Although most mitochondrial proteins can be imported into mitochondria as mature protein products post-translationally (3); the existence of cytosolic ribosomes associated with mitochondria in yeast was reported in early work by Kellems & Butow (4, 5) and more recently visualized in some detail (6). It is also well established that the mRNA of many nuclear encoded mitochondrial proteins localize primarily to the vicinity of mitochondria (7, 8, 9, 10), and for some particular genes (e.g. the F_1_-ATPase subunit ATP2) mRNA mitochondrial localization has been linked to mitochondrial function (11). Thus it is currently understood that both co-translational and post-translational mitochondrial import occurs in a wide range of eukaryotic organisms (12); although the exact delineation of the role of these two import modes with partially overlapping substrates remains unclear.

Previous studies have revealed several proteins whose deletion in yeast leads to significantly reduced mRNA mitochondrial localization. Deletion of the nascent polypeptide-associated complex NAC, a peripheral component of ribosomes which interacts with nascent chains as they leave the ribosome, leads to a significant reduction of mitochondrially associated ribosomes (and presumably mRNA), mitochondrial defects and reduced import of some mitochondrial proteins (13). Saint-Georges et al. (9) showed that the Pumilio family RNA binding protein Puf3p promotes the mitochondrial localization of the mRNA of 256 genes, highly enriched in assembly factors of respiratory chain complexes and the mitochondrial translation machinery. Tom20, a receptor subunit of the mitochondria outer membrane translocon (TOM complex) mediates mRNA mitochondrial localization in a translation-dependent manner and Tom20ΔPuf3Δ double knockouts have growth defects under conditions where fully functional mitochondria are required (14). Ssa1 is a general protein chaperone whose deletion reduces mRNA mitochondrial localization, especially for genes encoding hydrophobic proteins, in a way which is independent (additive) to protein N-terminal mitochondrial targeting signals, but dependent on Tom70 — another receptor subunit of the TOM complex (15). Finally, Lesnik et al. (12) identified the outer membrane protein OM14 as a receptor for nascent polypeptide-associated complex (NAC complex) and demonstrated its involvement in mRNA localization and co-translational protein import into mitochondria. Note that, except Puf3p, these proteins all affect mRNA mitochondrial localization via the nascent peptide chain or associated protein complexes.

A few features have been identified as characteristic of (various subsets of) mRNA which localize to the mitochondrial surface. Anderson & Parker (16) identified a 10-nucleotide motif enriched in mRNA encoding mitochondrially imported proteins, which was subsequently demonstrated to be the binding target of Puf3p (17) and to be specific to mitochondrially localized mRNA. Marc et al. (7) noted that mRNA encoding proteins with detectable bacterial homologs have a greater tendency to localize to mitochondria. The N-terminal mitochondrial targeting signals found in most proteins imported to the mitochondria matrix or inner membrane, have been shown experimentally to mediate the mitochondrial localization of their mRNA (15, 18). The mRNA of proteins involved in respiratory chain assembly have been noted to localize to yeast mitochondria in a Puf3p dependent manner (19). Similarly, in plants, the mRNA of respiratory, mitoribosome, and Krebs cycle genes exhibit mitochondrial localization (20). Other studies have pointed out that the mRNA of mitochondrial inner membrane proteins have a higher tendency to localize to the mitochondrial surface (10). Finally, protein length shows a strikingly strong positive correlation with the mitochondrial localization of its encoding mRNA (8).

Although the previously described mRNA localization studies have generated substantial experimental data for yeast (7, 8, 9, 10, 14), a combined analysis of these datasets has not been reported. Here, we perform an initial general analysis confirming previously noted trends; and then home in on previously unexplored mRNA sequence features related to translation initiation. Interestingly, all of the translation initiation related features we investigated indicated that rapid translation initiation corresponds with a *lower* degree of mRNA mitochondrial localization. This is surprising given the evidence listed above implicating the product peptide chain as a major determinant of mRNA mitochondrial localization.

To explain this seemingly paradoxical observation, we hypothesize that rapid initiation of translation of newly minted mRNA emerging from the nucleus leads to reduced mobility of the mRNA, decreasing the likelihood that those mRNA reach mitochondria; and we present a simple simulation to explore this idea. Intriguingly, short mRNAs are generally considered to tend to initiate translation rapidly, leading to the possibility that rapid translation initiation may be one reason short mRNAs tend not to localize at mitochondria.

## MATERIALS AND METHODS

### Mitochondrial mRNA Localization Value Datasets

We began our study with four mRNA mitochondrial localization datasets obtained with microarray technology (7, 8, 9, 14), which we label as MLR2002, MLR2003, MLR2008 and MLR2010 according to their year of publication. Each of these studies provide gene specific *Mitochondrial Localization of mRNA* values (MLR values), which reflect the degree to which mRNA molecules of that gene localize to mitochondria (i.e. a high MLR value indicates a tendency to localize at the surface of mitochondria, and a low MLR value a tendency to localize elsewhere in the cytosol). Restricting attention to genes whose protein products are imported into mitochondria; MLR2002, MLR2003, MLR2008, MLR2010 contain 344, 588, 564 and 280 genes respectively. The genes covered largely overlap, with the four datasets containing only (4, 25, 21, and 0) unique genes respectively. In general the agreement of MLR values between datasets is reasonably good, with pairwise correlation coefficients across experiments ranging from 0.45 to 0.79 (Supplementary Figure S3).

After our preliminary analysis with these datasets, we further corroborated our observations using two ribosome profiling based datasets from Williams et al. (10); which we denote as CHX+ and CHX-respectively depending on whether the translation elongation inhibitor cycloheximide (CHX) was added prior to measurement. CHX+ and CHX-contain 654 genes and correlate with a Pearson correlation coefficient of *r* = 0.72. Since the MLR values between the four microarray datasets displayed strong correlation (Supplementary Figure S3) and showed similar trends when plotted against features such as the ORF length (Supplementary Figure S1), we combined the four microarray based MLR datasets into a single dataset, which we denote as MLR4. To compute MLR4: for each dataset we linearly converted the MLR values to a scale between 0 and 1, and then for each gene averaged the available MLR values to produce a merged dataset. MLR4 contains 643 genes and has a correlation coefficient of *r* = 0.66 with CHX+ and r = 0.53 with CHX-, respectively.

### Multivariate Regression Analysis

To investigate the factors involved in mRNA localization, we compiled data for various features of mRNA and their protein products. As described in the supplementary material text, we used both feed forward neural networks and linear regression to train predictors of MLR value from these features. Table 1 lists the features we used for regression, along with ribosome profiling data based estimates of translation initiation probabilities (21) discussed later.

**Table 1.** Correlation between mRNA (or protein product) features and MLR values averaged over the four microarray datasets. *n* denotes the number of genes for which a value for the given feature was available, for each respective MLR dataset; *r* and *ρ* are the Pearson and Spearman rank correlation respectively. Features with unexpectedly negative correlation with MLR are shown in gray.

| Feature | avg. $n$ | avg. $r$ | avg. $\rho$ | Source |
| --- | --- | --- | --- | --- |
| ORF length | 446.5 | 0.42 | 0.55 | Uniprot |
| ORF length (cube root) | 446.5 | 0.53 | 0.55 | Uniprot |
| ORF length (log) | 446.5 | 0.56 | 0.55 | Uniprot |
| Translation initiation probability | 428.5 | -0.39 | -0.33 | Weinberg et al. (21) |
| Ribosome occupancy | 414.75 | -0.29 | -0.29 | Arava et al. (25) |
| Ribosome density | 414.75 | -0.36 | -0.35 | Arava et al. (25) |
| Ribosome density | 410.5 | -0.12 | -0.13 | as listed by Siwiak et al. (24) |
| Ribosome occupancy × Arava Ribosome density | 414.75 | -0.39 | -0.36 | Density also from Arava et al. (25) |
| Ribosome occupancy × Siwiak Ribosome density | 383.5 | -0.15 | -0.13 | Density as listed by Siwiak et al. (24) |
| Number of ribosomes | 410.5 | 0.18 | 0.26 | as listed by Siwiak et al. (24) |
| Transcript abundance | 400.75 | -0.18 | -0.17 | Nagalakshmi et al. (22) |
| tRNA adaptation index (tAI) | 446.5 | -0.04 | -0.00 | Reis et al. (26) |
| 5'UTR length | 374.5 | 0.07 | 0.07 | Nagalakshmi et al. (22) |
| 3'UTR length | 374.5 | 0.02 | -0.00 | Nagalakshmi et al. (22) |
| mRNA half-life | 393.25 | -0.03 | -0.03 | Miller et al. (23) |
| mRNA synthesis rate | 393.25 | -0.19 | -0.14 | Miller et al. (23) |
| Disorder of N-terminal 50 AA residues | 446.5 | 0.25 | 0.25 | Prediction by DISOPRED (27) |
| Hydrophobicity | 446.5 | 0.03 | 0.02 | Calculation by Kyte & Doolittle index |
| Has MTS (by experiment) | 249.25 | 0.28 | 0.28 | Vögtle et al. (28) |
| Has MTS (by prediction) | 446.5 | 0.31 | 0.30 | Emanuelsson et al. (29) |
| Prokaryote homolog search log E-value | 431.5 | -0.28 | -0.33 | SSEARCH (Pearson (30)) |
| Log protein expression level | 351 | 0.04 | 0.08 | Ghaemmaghami et al. (31) |
| Protein half-life | 322 | -0.05 | -0.13 | Belle et al. (32) |
| Puf3 binding | 109.5 | 0.05 | 0.05 | Gerber et al. (17) |
| ΔSsa1 effect | 186.5 | 0.49 | 0.53 | Eliyahu et al. (15) |
| ΔTom20 effect | 255 | 0.23 | 0.21 | Eliyahu et al. (14) |

## RESULTS

### Multivariate Regression

The main conclusion we drew from the regression analysis is that the ORF length is statistically the dominant factor. The logarithm of the ORF length has a stronger linear correlation with MLR value than the ORF length itself, presumably because the logarithm is used in computing the MLR values from experimental measurement. For brevity, we will sometimes omit the “logarithm of” when discussing this feature. As we discuss in the following section, ribosome occupancy (defined as the fraction of mRNA molecules of a given gene occupied by one or more ribosomes), ribosome density and translation initiation probability have a fairly strong correlation with MLR, but surprisingly it is negative. Other features in Table 1 have some relationship with MLR, for example there is a statistically significant tendency for genes encoding proteins with short half-lives to localize to mitochondria (33), but the overall correlation between those features and MLR is weak.

Not only does ORF length predict MLR values much better than the other features (Supplementary Figure S1), combining ORF length with the other features only marginally improves the prediction (Supplementary Table S1) over using just ORF length. In two out of four datasets tested, an N-terminal mitochondrial matrix targeting signal (MTS) statistically significantly improves prediction of MLR when combined with ORF length; presumably due to its contribution to sustained anchoring of mRNA at the mitochondrial surface via translocation of the N-terminus of nascent polypeptide chains into the mitochondria. In one out of four datasets tested, the ΔSsa1 effect feature can be combined with ORF length to improve prediction of MLR. Actually ΔSsa1 is not a feature of the mRNA per se, but rather a measurement of how much the MLR of that gene changed when overexpressing Ssa1 (15). Thus ΔSsa1 is derived from MLR measurements, and therefore it is not surprising that it correlates with MLR values from other experiments as well. We included it in our analysis in case we might find an interesting combination of ΔSsa1 with other mRNA features which together could substantially increase the power to predict MLR. However, although statistically significant in some cases, the improvement in MLR prediction achieved by considering these features in addition to ORF length is modest; and overall the ORF length can be said to be the statistically dominant feature. These results confirm and strengthen earlier observations by Sylvestre et al. (8), who reported a strong correlation between the logarithm of the ORF length and the degree of MLR, and also concur with the aforementioned ribosome profiling experiment by Williams et al. (10) in which they found that shorter ribosome-nascent chains are seldom observed near mitochondria.

#### Anti-Correlation Between MLR and Ribosome Occupancy and Density

Contrary to our prior expectations, MLR displayed a negative correlation with both ribosome occupancy and ribosome density. This surprised us because an explanation of the length-MLR correlation proposed by Sylvestre et al. (8), based on a model of George et al. (13), invokes ribosomes connected to the mitochondrial surface via the nascent-polypeptide-associated complex (NAC). The argument consists of three logical steps: 1. ribosomes help anchor mRNA to the mitochondria, 2. longer mRNA have more ribosomes, and therefore — 3. longer mRNA localize to mitochondria.

Given this background, we initially found the negative correlation between ribosome occupancy and MLR the most surprising, and therefore we double-checked by computing the correlation between ribosome occupancy and MLR4 (i.e. correlation with the average MLR over datasets instead of the average of the correlation over datasets shown in Table 1). For the 543 genes with data available for ribosome occupancy, number of ribosomes and MLR4 plotted in Figure 1, the Pearson correlation of ribosome occupancy with MLR4 is −0.378, with a *p*-value < 2.2 × 10^-16^ — the limit of what the *t*-test based R language function we used can compute.

**Figure 1.**
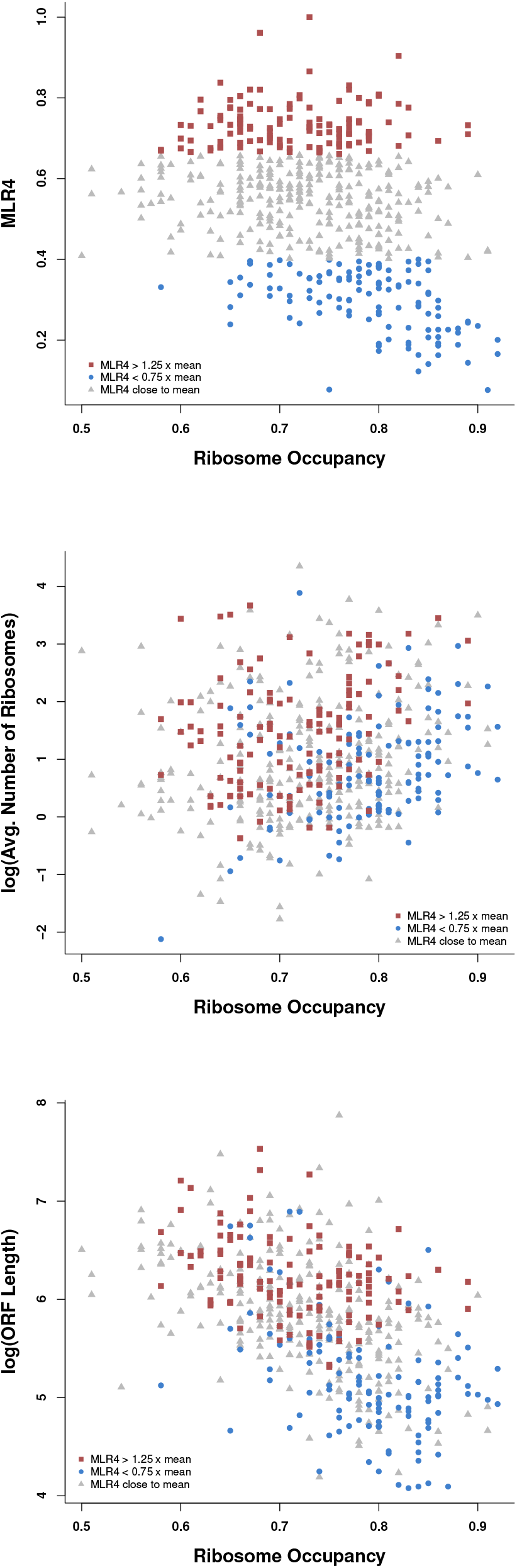
Ribosome occupancy of yeast mRNAs coding for mitochondrially imported proteins plotted against: MLR4 (top); the natural log of their average number of ribosomes (middle); and the natural log of the length of their product in codons (bottom). Points are colored based on their MLR4 value.

Admittedly not all ribosome related features anti-correlate with MLR. As can be seen in Table 1, the average number of ribosomes (using data listed by Siwiak et al. (24), from the ribosomal profiling experiments of Ingolia et al. (34)) correlates moderately positively with MLR (average correlation: 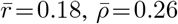), with some variation between datasets (Supplementary Figure S4). This positive correlation is consistent with the notion that ribosomes are relevant to MLR, as would be expected by the above model and the fact that ribosomes associate via NAC with the mitochondrial outer membrane protein OM14(12). Nevertheless the relatively modest correlation between MLR and average number of ribosomes (vis *á vis* log ORF length) and the negative correlation with ribosome occupancy and density strongly suggests there is more to the relationship between MLR and ORF length than just ribosome mediated anchoring. Interestingly, ribosome occupancy and average number of ribosomes together do complement each other well in separating mRNAs with a high or low degree of MLR — a point we revisit later.

#### Translation Initiation Deserves a Closer Look

The remaining feature to be discussed is translation initiation. As seen in Table 1, the translation initiation probabilities estimated by Weinberg et al. (21) show a clear negative correlation with MLR (average correlation: 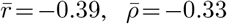). Interestingly the negative correlation between MLR and ribosome occupancy could potentially be explained by a negative correlation between MLR and rapid translation initiation. Ribosome occupancy depends on the rate of translation initiation as well as the elongation and termination times. Arava et al. (25) observed however that the ribosome density for the vast majority of mRNAs was much lower than the maximal capacity, suggesting that initiation is generally the translation limiting factor. Indeed, if occupancy were strongly dependent on the elongation rate one would expect that longer mRNA should have high ribosome occupancy, since if the time needed for a ribosome to clear long mRNA far exceeds the mean time for a new translation initiation, long mRNAs would continuously be in a polysome state; but in fact mRNA with longer ORFs have *lower* ribosome occupancy (Figure 1).

Another reason to take a closer look at translation initiation rate is that it is believed to correlate strongly with ORF length — the strongest single predictor of MLR identified so far. Several (but not all (24, 35)) studies in yeast have shown a strong negative correlation between the ORF length and computationally derived translation initiation likelihoods (36, 37), and in experiments both translation efficiency (21) and estimated translation rates (25) have shown the same negative correlation with length. Experiments in mammalian cells also support the assertion, with a strong negative correlation between ORF length and translation rates reported in two different cell types (38). Thus we adopt the working hypothesis that shorter mRNAs tend to initiate translation more rapidly. The above observations prompted us to further explore the relationship between MLR and translation initiation by investigating the correlation between MLR and mRNA sequence features (e.g. Kozak sequences) known to promote rapid translation initiation.

### Translation Initiation Promoting Sequence Features anti-correlate with MLR

#### Kozak Sequences

We investigated the relationship between MLR and translation initiation promoting Kozak sequences. In yeast Kozak sequences are characterized by a preponderance of adenine immediately upstream of the initiator `AUG` codon, especially in the −3, and to a lesser extent −1 position; and `UCU` or a related codon in the +4 to +6 position (39, 40).

First we investigated the correlation between MLR and the frequency of nucleotides in the −1 to −10 region for the 494 mRNA sequences with 5′ UTRs of length > 10. Overall, there is high frequency of `A`s; a known preference in yeast 5′ UTRs (42), with the mean MLR value of mRNAs with an `A` falling below the overall average in every position except −10 (Figure 2). Similar results can be seen in the CHX-dataset where all `A`s were associated with lower than average MLR values except at position −4 (Supplementary Figure S5). The CHX+ data was less clear with 6 `A`s showing below average MLR values. Conversely, `G`s and `C`s tended to be associated with higher MLR values than `A`s in all the datasets (statistically significantly so in MLR4: *p* = 0.0098 and CHX-: *p* = 0.020, Wilcoxon *T* test). Since `G+C` rich pair stem-loop structures have been found to inhibit translation initiation in yeast (43, 44), this result may indicate that mitochondrially localizing mRNAs tend to have more secondary structure and consequently lower translation initiation rates (a question we address more directly later).

**Figure 2.**
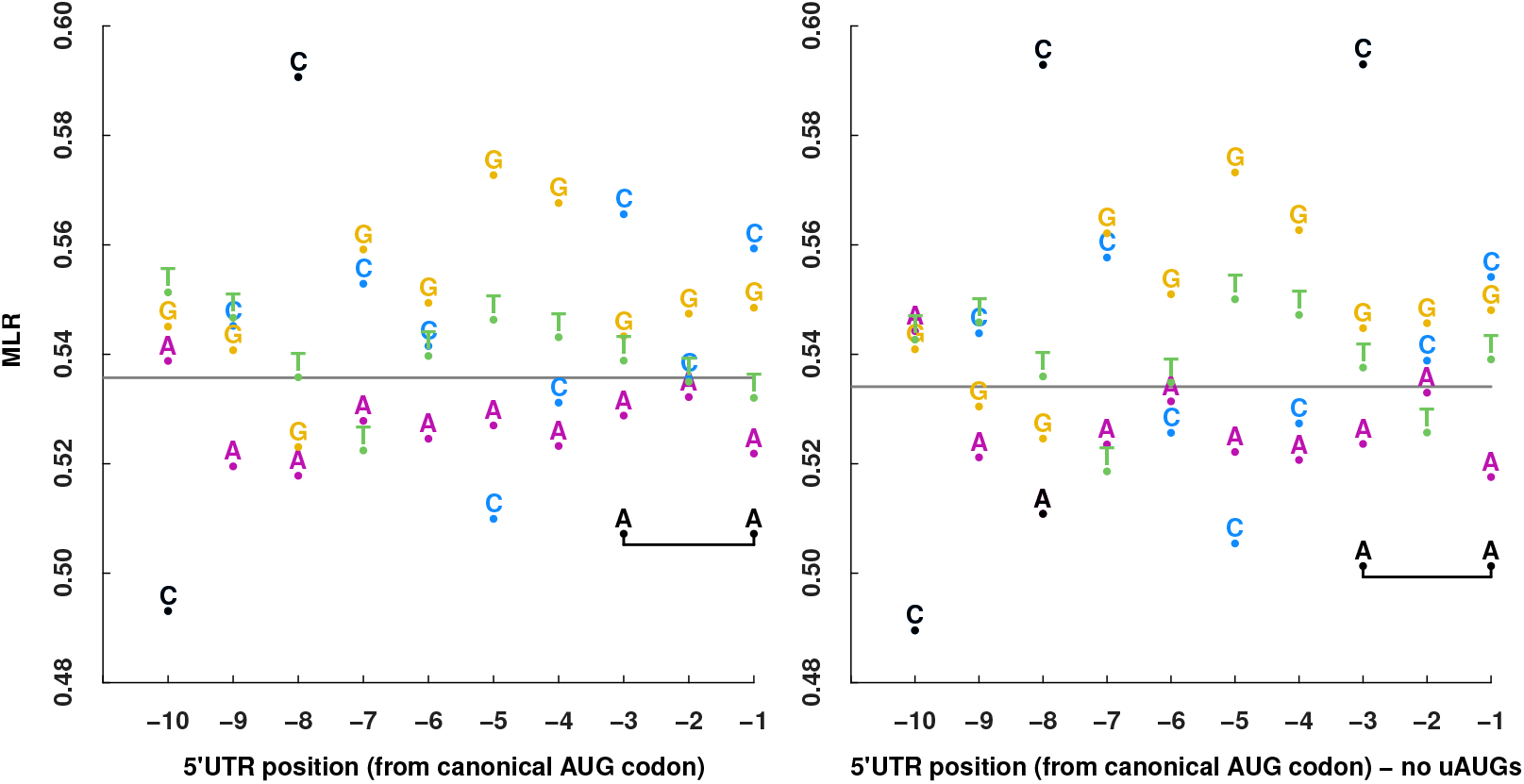
The average MLR4 value for mRNA with an `A, C, G`, or `T` respectively, at the −10 to −1 positions upstream of the translation start codon for all mRNAs (left) and those with no upstream `AUG`s (right) are shown. Black color indicate position/base combinations whose average MLR4 exhibit relatively large statistical deviations from the overall average MLR4 value (horizontal bar). Plot style based on Dvir et al. (41).

We also observed some correlation between MLR and particular bases at particular positions upstream of the starting `AUG` (Figure 2). In particular, the mean MLR value of the 132 mRNAs with an `A` at both −3 and −1 (`A.AAUG`s) is statistically significantly lower than the overall average (*p* = 0.019, independent *t*-test), without multiple testing correction since we performed the test based on prior knowledge that A’s at the −3 and −1 positions are especially important for efficient translation initiation(39, 41). `A.AAUG`s were also associated with lower MLR values in the CHX+ and CHX-datasets, statistically significantly so in the CHX-dataset (*p* = 0.0018) (Supplementary Figure S5). Before correction for multiple testing, several individual nucleotide/position pairs appeared potentially significant: a `C` at −10 and −5 associated with lower MLR values (*n* = 78, *p* = 0.022 and *n* =107, *p* = 0.033 respectively), and a `C` at −8 (*n* = 82, *p* = 0.0005) and a `G` at −5 (*n* = 65, *p* = 0.038) associated with higher than average MLR values. However, after applying Bonferroni correction, only the C at −8 remained statistically significant (corrected p = 40*0.0005 = 0.02). As far as we are aware, a `C` at −8 has not been mentioned previously in any context (e.g. translation efficiency) in the literature, so it is difficult to interpret this result — except to note that its correlation with MLR is unlikely to be due to chance.

To further clarify the correlation between Kozak sequences and MLR, we repeated this analysis using only the 405 (out of 494) mRNA sequences with no upstream `AUG`s (which could potentially complicate the picture by competing for translation initiation). Comparing the left and right panels of Figure 2, one can see that the removal of the 89 mRNA with upstream `AUG`s tended to increase the differences in mean MLR value, especially for an `A` at −8 (which becomes statistically significant prior to Bonferroni correction) and increases the significance of the low mean MLR value observed in mRNAs with `A.AAUG`s (*n* =119, *p* = 0.010).

As summarized in Table 2, we also investigated the relationship between MLR and the second codon. In early work, Hamilton et al. (39) noted that `UCU` is the most common second codon in a collection of 93 yeast genes and especially common in highly expressed genes. In a more comprehensive comparison, Gingold and Pilpel (40), analyzing the translation profile data of Arava et al. (25), similarly demonstrated enrichment of `UCU` and other second codons ending in `CU` in highly efficiently translated genes. Interestingly, for each of the three sets of MLR measurements we analyzed, the MLR values of mRNA with `UCU` as their second codon was lower than mRNA with other codons (two-tailed, two-sample *t*-test; p-values; MLR4:9.7 × 10^-4^, CHX-:0.12, CHX+:0.036). This trend was even more pronounced when comparing mRNA with codons matching `XCU` versus the others (*p*-values; MLR4:3.8 × 10^-5^, CHX-:0.019, CHX+:0.0025). Note that these trends do not seem to be a by-product of coding for serine, as the average MLR values of the non-`UCU` serine codons are similar to the overall average (Table 2).

**Table 2.** Average MLR correlates with second codon. The number of genes n in the MLR dataset matching each pattern (“!” indicates negation) is shown followed by their mean values of MLR4, CHX+ and CHX-. The numbers after “±” are the standard error of mean estimation (SEM). Rows corresponding to translation initiation promoting second codons are colored gray.

| pattern | n | MLR4 | CHX+ | CHX- |
| --- | --- | --- | --- | --- |
| !XCU | 580 | .536±.007 | 1.283±.059 | .711±.034 |
| XCU | 73 | .449±.019 | 0.720±.171 | .491±.085 |
| !UCU | 606 | .532±.007 | 1.254±.058 | .699±.033 |
| UCU | 47 | .447±.024 | 0.779±.213 | .522±.108 |
| UCV | 59 | .526±.020 | 1.046±.184 | .695±.098 |

#### Upstream AUGs

Ribosomes typically initiate translation using a scanning model that favors the `AUG` nearest to the 5′ end (45, 46, 47, 48). Thus upstream ORFs (uORFs, defined as a pair of equal-frame start and stop codons in the 5′ UTR) can compete with the canonical start `AUG` for translation initiation, leading to a substantial reduction in protein production (49, 50, 51). This prompted us to compare the MLR values of mRNA with (*n* = 55) and without (*n* = 435) uORFs (Supplementary Figure S7). We observe that mRNAs with uORFs have higher MLR values than mRNAs without uORFs, almost significantly so in some cases (*p* = 0.095 and 0.066 for the CHX+ and CHX-datasets respectively, independent *t*-test). This results differs from a study of plant cells, in which uORFs were associated with reduced mRNA mitochondrial localization (20).

In the previous section we reported that the upstream context of the canonical `AUG` codons correlated with MLR. Therefore, we performed a similar analysis on uORF `AUG`s; computing the average MLR value for such `AUG`s in different contexts. To simplify interpretation of the results, we initially only considered mRNA with a single uORF (more precisely, a single uORF after discarding any short uORFs nested inside a longer ORF in the same frame) yielding a sample size of *n* = 29 for MLR4 and *n* = 30 for CHX+ and CHX-. As shown in Figure 3 (MLR4 results) and Supplementary Figure S8 (CHX+ and CHX-results), the correlation between MLR and context for these u`AUG`s differ markedly from the correlation observed with canonical start `AUG`s. In particular, `A`s upstream of uORF `AUG`s tend to have higher instead of lower than average MLR values. Comparing the canonical start `AUG` and u`AUG A` counts below and above the MLR mean, we found a significant difference in the MLR4 and CHX-datasets (*p* = 0.020 and *p* = 0.0039, Fisher’s exact test).

**Figure 3.**
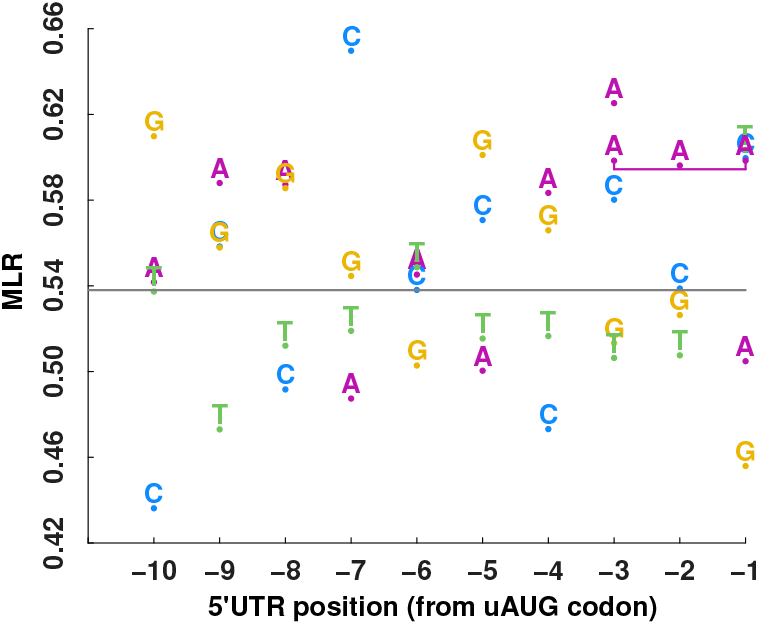
Similar plot to figure 2, but showing the context of the upstream `AUG` of single uOrF mRNAs.

Interestingly, the increased MLR value was especially noticeable for the handful of single uORF containing mRNAs with an optimal u`AUG` context of `A.AAUG`, reaching statistical significance for the CHX-set (*p* = 0.045, Wilcoxon *T* test) despite the small sample size (*n* = 4). Consistently, when allowing multiple uORFs, the average MLR values of u`A.AAUG`s (*n* =11) were higher (ΔMLR = 0.04, 0.35, 0.31, for MLR4, CHX+ and CHX-respectively) than non-`A.A` u`AUG`s (MLR4: *n* = 69, CHX+-: *n* = 71), significantly so for CHX- (*p* = 0.0052, Wilcoxon *T* test).

#### 5′ *UTR and* 3′ *UTR Nucleotide Distributions*

We compared the nucleotide distribution of MLRs between the 5′ UTR and 3′ UTR region in positions 0 to 50 from each respective UTR end as the 5′ UTR appeared to show greater variance (Figure 4). Because the sample size at each position in the 5′ UTR and 3′ UTR region differs, we only included nucleotides with sufficient sample size at each position (see supplementary material text for details). The 5′ UTR MLR nucleotide distribution variance was greater than the 3’ UTR distribution in all datasets (higher in the 5′ UTR by 60%, 51% and 74% in the MLR4, CHX+ and CHX-datasets respectively), significantly so in the case of the MLR4 and CHX+ datasets (F-test; MLR4: *p* = 0.0082; CHX+: *p* = 0.00049, CHX-: *p* = 0.11, Supplementary Figure S9). Although the 3’ UTR can have an impact on translation initiation in some cases (55), the 5’ UTR affects translation efficiency more strongly (56). Thus the larger association between particular nucleotides and MLR observed in the 5’ UTR region again hints at a possible role for translation initiation rate in mRNA mitochondrial localization.

**Figure 4.**
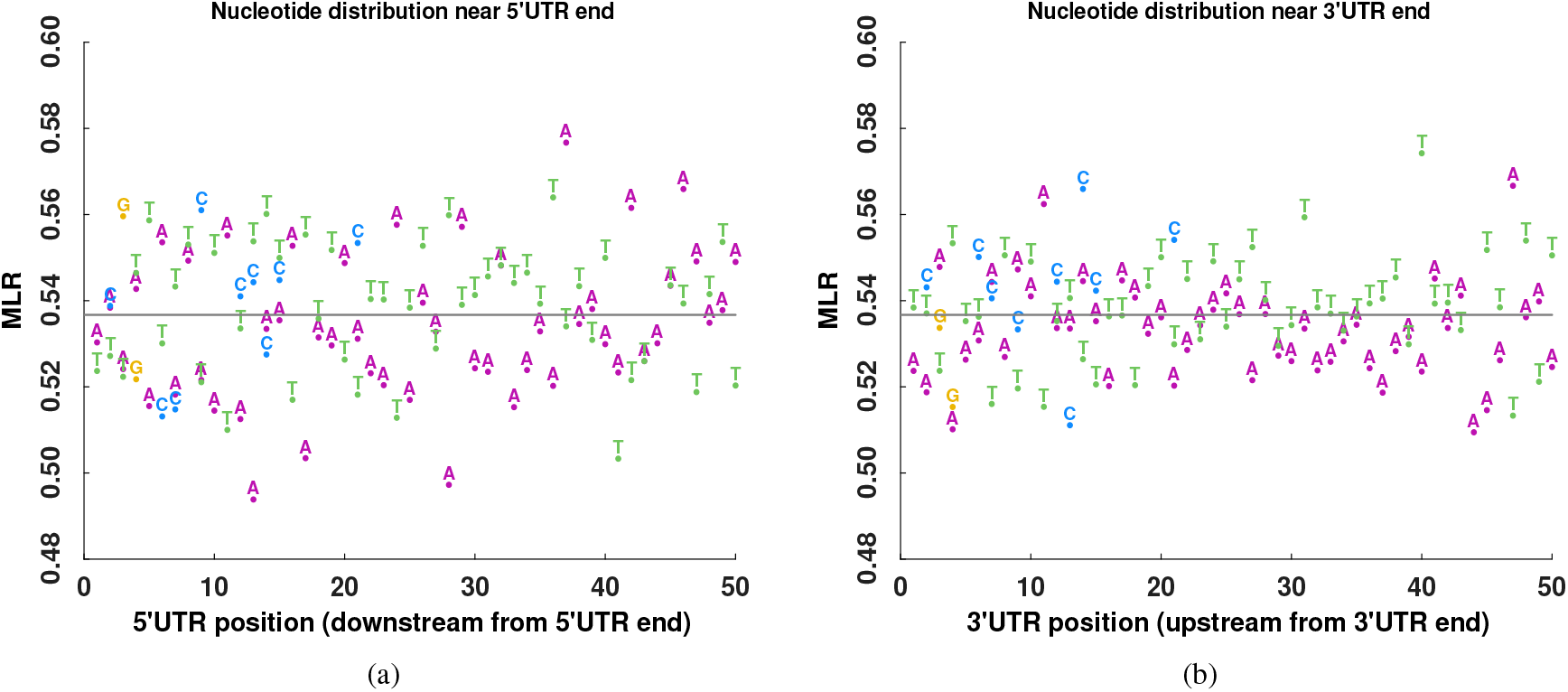
The average MLR value mRNA with an `A, C, G`, or `T` nucleotide at positions downstream from the 5′ UTR (a) and 3′ UTR (b) ends is shown. The gray horizontal line indicates the average MLR value across all mRNA. Only nucleotides with sufficient counts are shown (see supplemental text for details).

#### Secondary Structure

Using calculated minimum free energy (MFE), Dvir et al. (41) found that thermodynamically stable structured mRNA (lower MFE) have relatively low protein abundance. Similar results have been reported in small scale experiments, where early studies found that secondary structure could affect ribosomal initiation in the herpes thymidine kinase (57) and rat preproinsulin (58) genes, followed by results in yeast that likewise reported secondary structure inhibiting translation initiation (43, 52, 59, 60, 61). This prompted us to compare MFE and MLR values for our set of mRNA. Following Dvir et al. (41), we used zipfold (62) to calculate MFE values for the range of −15 to +50 nucleotides relative to the start codon. As shown in Figure 5 (MLR4 data) and Supplementary Figure S10 (CHX+ and CHX-data), we observed a highly statistically significant negative correlation between MLR and MFE for all three datasets; again consistent with the conclusion that rapid translation initiation (afforded by unstructured mRNA with high MFE) is associated with a small degree of mRNA mitochondrial localization.

**Figure 5.**
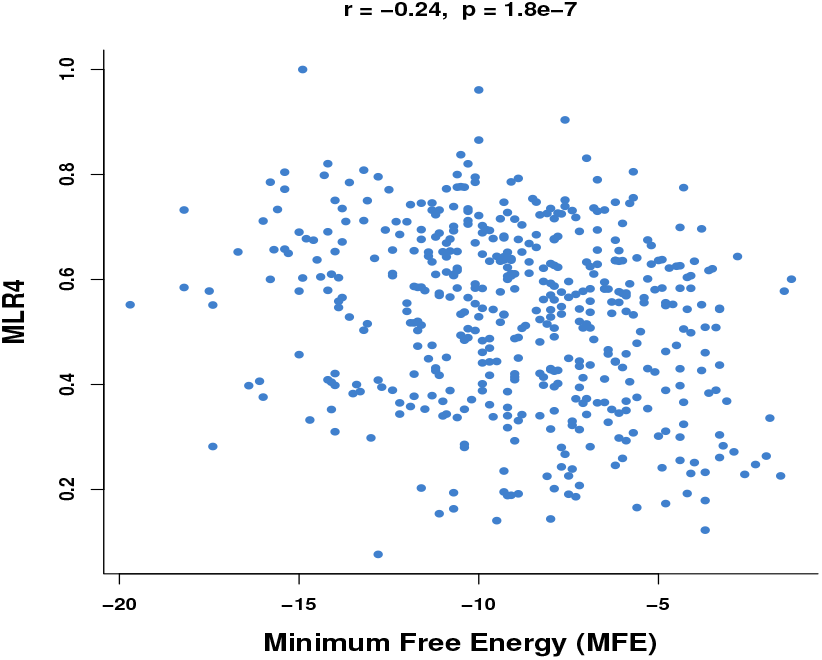
Scatter plot of the calculated minimum free energy of the −15 to +50 region of mRNA versus their MLR4 values. *r* and *p* are the values of the Pearson correlation coefficient and its statistical significance by *t*-test against a null hypothesis of no correlation.

### A New Model: Diffuse then Drop Anchor

The translation initiation related sequence features we investigated all point to slow translation initiation correlating with a high tendency for mRNA to localize to mitochondria. This might seem surprising, since most mechanisms proposed for anchoring mRNA to the mitochondria involve the nascent polypeptide chain or associated proteins — and therefore require translation. In this section we elaborate on a new model which attempts to resolve this apparent contradiction. Our model follows the “diffuse and anchor” mechanism proposed for mRNA localization in some other contexts, such as the localization of *nanos* mRNA to the *D. melanogaster* pole plasm (63), but more closely follows the ship-and-anchor analogy by assuming that the anchor (the nascent polypeptide chain and associated proteins) will slow down the ship if dropped prematurely (i.e. translation initiates far from a mitochondrial surface). Thus our model has two main assumptions: first, that mRNA molecules become less mobile after they initiate translation and therefore have a reduced chance to encounter the surface of a mitochondrion; and second, that mobile mRNA molecules (at least those coding for mitochondrial proteins) are more likely to initiate translation when in the vicinity of mitochondria. Under these assumptions an mRNA molecule that defers initiating translation until encountering the mitochondrial surface will more likely localize there than an mRNA molecule which initiates translation soon after nuclear egress.

Why might mRNA molecules become less mobile once they initiate translation? Studies on mammalian cells indicate that translating ribosomes are often associated with the cytoskeleton (64, 65, 66) which plays an active role in regulating translation (67, 68), but also presumably reduces the ability of the translating mRNA molecules to diffuse freely (69, 70, 71). In addition to diffusion, the possibility of (cytoskeleton mediated) active transport enabled via recognition of *cis* “zipcode” sequences must also be considered, as many examples of mRNA localization via this mechanism are known (72). However mRNA translation is typically silenced during transport (73, 74), which is at least qualitatively consistent with our assumption that translation initiation reduces mRNA mobility. In any case diffusion (thermal or ‘active’ (75)) appears to be centrally important for the movement of most mRNA. Indeed, visualizing the movement of single molecules of reporter mRNA in COS cells, Fusco et al. (76) found only 20% and 2-4% of the molecules exhibited directed motion for *β*-actin zipcode-containing and zipcode-free mRNAs respectively. These studies were mainly done on mammalian cells, but since the cytoskeleton in yeast cells is denser than in mammals (77), one would expect association with the cytoskeleton via translating ribosomes to hinder diffusion at least as much in yeast as in mammalian cells.

Our second assumption is that mobile mRNA molecules are more likely to initiate translation at or near mitochondria. This statement follows to some degree simply from the fact that encountering mitochondria is a prerequisite for initiating translation there. Indeed, the mitochondrial surface is rich in ribosomes (4, 5), a certain fraction of which might be available for engaging new mRNA partners at any given time. However, this raises the question of why mobile mRNA molecules do not initiate translation on the even more numerous ribosomes distant from mitochondria. To explain the high level (in many cases >50%) of mitochondrial localization attained by the mRNA of many genes, the existence of a mitochondria-specific mechanism must be assumed.

#### Our Model Can Explain Ribosome Occupancy Trends

Although our model is a simplification which does not attempt to explain all observations reported in the literature, we investigated how well it can explain trends found in the MLR datasets. In particular, we note that our model can account for the mixed results between the number of ribosomes (positive correlation with MLR) and ribosome occupancy. Indeed, despite a lower average ribosome occupancy, high MLR value genes tend to have a higher average number of ribosomes (Figure 1, Supplementary Figure S11). Our model explains this as the result of two competing effects: on the one hand more ribosomes (and more nascent polypeptides) help anchor mRNA to the mitochondria as in earlier models (8, 13), but on the other hand mRNA carrying no ribosomes are more mobile and therefore more likely to encounter mitochondria. Thus our model can be viewed as an extension of previous explanations, adding consideration of the effect of translation initiation on mRNA mobility.

#### Exploratory Simulation

To explore our model we implemented a discrete time mRNA diffusion simulation which incorporates the two main assumptions of our model: reduced mobility of translating mRNA and an increase in the probability of translation initiation near the mitochondria. The simulation inputs gene specific translation initiation rates and outputs simulated MLR values. We used translation initiation rates from Shah et al. (37), and for each gene we simulated the (translation state dependent) diffusion and entrapment of an mRNA molecule multiple times to obtain a simulated MLR value for that gene.

The simulation is described in detail in the supplementary material. Briefly, we modeled a cell as a sphere of radius 2.5*μ*m containing 1 to 5 spherical mitochondria with radii of 0.25-0.5*μ*m. An mRNA molecule starts in an nontranslating state, drifting unhindered in the cytosol. As the simulation progresses the molecule may become entrapped or encounter a mitochondrion before or after initiating translation (elongation and termination of translation are not modeled), or possibly wander through the cytosol until the end of the simulation. The probability of entrapment is increased upon initiating translation and the probability of translation initiation is increased upon encountering a mitochondrion. For the diffusion coefficients of translating and non-translating mRNAs, the main simulation adapts molecule size based estimates by (76); while a companion follow-up study explores a range of diffusion coefficients.

Parameters used to explore our hypothesis are *x* (the ratio of the probability of an mRNA molecule becoming entrapped in any given time step before *vs.* after initiating translation) and the mRNA Diffusion coefficients of translating and nontranslating mRNA (see supplementary material for details). Encouragingly, the simulation approximately recreates the MLR values (*r* = 0.44) from Sylvestre et al. (8) and shows a relationship with ORF length similar to that of real MLR values (Figure 7) when *x* is set to 5 (and qualitatively similar for values from 5-20). As a control, when *x* is set to 1 (i.e. when translation initiation is assumed to have no effect on the probability of entrapment) and the same diffusion rate was used for translating and non-translating mRNA, then simulated MLR showed no correlation with real MLR values (*r* = 0.02).

**Figure 6.**
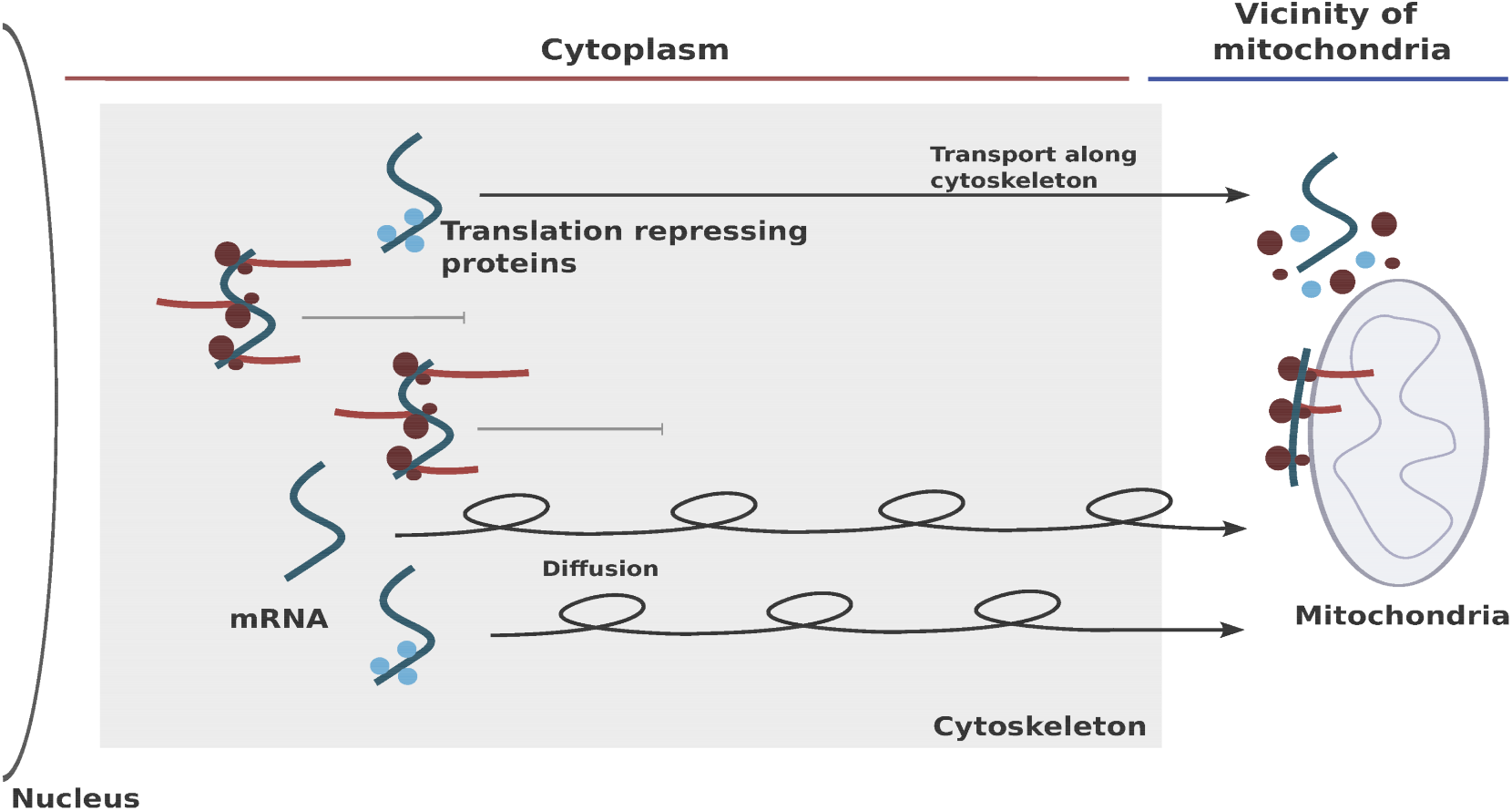
Model of mRNA localization to mitochondria. Short lines that end with a horizontal line depict the hindered movement of translating mRNA whose ribosomes and nascent peptides (marked by solid red circles on mRNA and protruding lines) may transiently or persistently become immobilized due to contact with the cytoskeleton. In contrast, mRNA with slow translation initiation (either inherit in the mRNA sequence itself, or mediated by translation repressing proteins shown as solid blue circles) are more likely to diffuse freely (loopy arrows) or actively move along the cytoskeleton (straight arrows). mRNA may initiate translation upon arrival at the mitochondrial surface, for example due to the disassociation of translation repressing proteins, and subsequently anchor there via their nascent polypeptide chain or associated proteins. (For simplicity free mRNAs are depicted as isolated molecules, even though they are typically packaged in ribonuclear protein particles).

**Figure 7:**
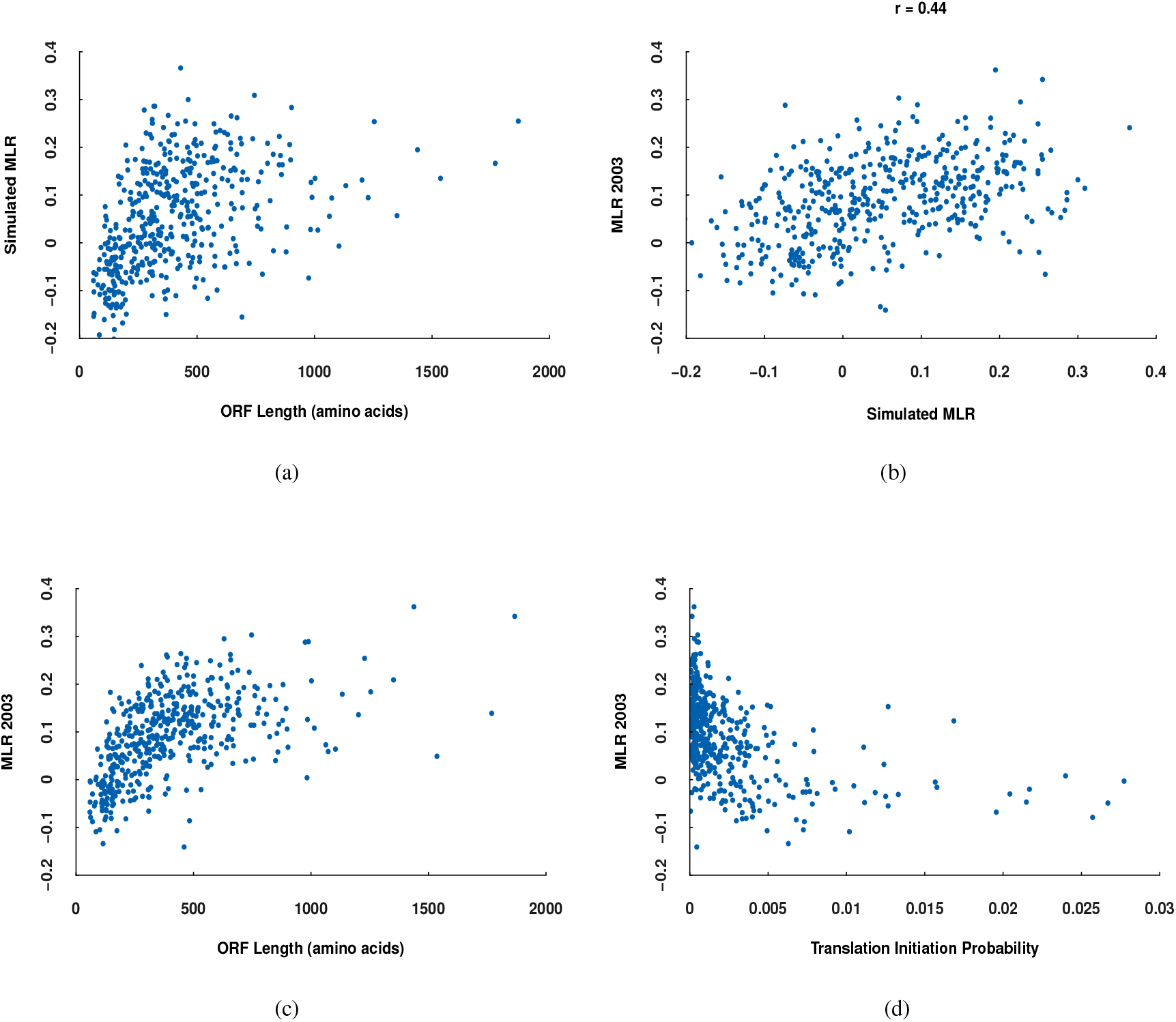
For each gene we computed a simulated MLR value: a) The relationship between simulated MLR values and the ORF length was similar to the real data shown in c). b) The simulated MLR values correlated well with the real MLR data. c) Comparison between the ORF length and MLR values from Sylvestre et al. (8). d) The translation initiation probabilities used as input for the simulation have a non-linear relationship with MLR distinct from the ORF length and the simulated MLR values.

We next wanted to ensure that the simulated MLR values were indeed dependent on the translation initiation rates, and not a byproduct of the simulation setup. We therefore ran the simulation with a randomly generated set based on translation initiation times (where a Gaussian distributed set was generated using the mean and variance from the Shah et al. (37) translation initiation times, such that the same number of samples was created as in the real data). As can be seen in Figure S13, random translation initiation rates fail to reproduce MLR values similar to actual MLR values.

For simplicity, in the simulations reported above we stipulated that mRNA above always anchor when reaching a mitochondrion. To explore the effect of varying the anchoring probability, we also performed simulations with probabilities ranging from 0.05 to 1.00. As shown in Table 3, the linear correlation between simulated MLR (*r*) and the Sylvestre et al. (8) measurements was not very sensitive to the value of the anchoring probability. This is as expected as the primary effect of lowering the anchoring probability is to reduce the fraction of mitochondrially anchored mRNA molecules for all genes; but the coefficient of correlation is only sensitive to gene-specific changes, or changes which are highly non-linear even after taking the logarithm of the number of molecules in each location in accordance with the formula for MLR (8). Note also that if an mRNA molecule reaches a mitochondrion once, it typically will hit that mitochondrion multiple times, giving it multiple chances to anchor (see follow-up study).

**Table 3.** For each anchoring probability ten simulations were performed. *r* in the second column denotes the average Pearson’s correlation across the simulations; while the remaining columns list the average, minimum and maximum MAE (Mean Absolute Error) values.

| Anchoring Probability | $r$ | Average MAE | Min. MAE | Max. MAE |
| --- | --- | --- | --- | --- |
| 0.05 | 0.331 | 0.083 | 0.081 | 0.088 |
| 0.10 | 0.386 | 0.079 | 0.077 | 0.082 |
| 0.15 | 0.390 | 0.080 | 0.078 | 0.081 |
| 0.20 | 0.413 | 0.085 | 0.081 | 0.089 |
| 0.25 | 0.418 | 0.083 | 0.081 | 0.085 |
| 0.30 | 0.433 | 0.083 | 0.080 | 0.087 |
| 0.35 | 0.444 | 0.089 | 0.083 | 0.094 |
| 0.40 | 0.433 | 0.088 | 0.082 | 0.092 |
| 0.45 | 0.427 | 0.087 | 0.083 | 0.090 |
| 0.50 | 0.438 | 0.096 | 0.092 | 0.100 |
| 0.55 | 0.443 | 0.085 | 0.082 | 0.087 |
| 0.60 | 0.430 | 0.086 | 0.083 | 0.088 |
| 0.65 | 0.420 | 0.086 | 0.083 | 0.090 |
| 0.70 | 0.427 | 0.086 | 0.082 | 0.090 |
| 0.75 | 0.437 | 0.092 | 0.087 | 0.095 |
| 0.80 | 0.436 | 0.084 | 0.081 | 0.087 |
| 0.85 | 0.426 | 0.085 | 0.081 | 0.088 |
| 0.90 | 0.428 | 0.091 | 0.086 | 0.096 |
| 0.95 | 0.441 | 0.090 | 0.086 | 0.095 |
| 1.00 | 0.439 | 0.091 | 0.088 | 0.093 |

## DISCUSSION

We started our study by re-examining several published MLR datasets, looking for trends which might have been overlooked by the original authors. That preliminary study reaffirmed the known correlation between MLR and ORF length, and in fact found it to be a statistically dominant factor which predicts MLR almost as well by itself as when combined with other features. Fortunately our preliminary study also provided hints that we should take a close look at the relationship between MLR and translation initiation — an insight which also led us to rethink previous explanations of the MLR-length correlation. In our investigation into the relationship between mRNA translation initiation rate and mRNA mitochondrial localization, we consistently found translation initiation promoting sequence features to correlate negatively with MLR; and conversely, that translation initiation inhibiting features correlate positively with MLR. These results suggest that rapid translation initiation rate may prevent mRNA mitochondrial localization (but see the discussion below). And, if the commonly made assertion that shorter mRNAs tend to initiate translation quickly is true, offer a new way to explain the observed MLR-length correlation.

We incorporated these observations into a new model of mitochondrial mRNA localization which extends a previously proposed model by adding the assumption that non-translating mRNAs are more mobile than translating mRNAs and explicitly assuming that the translation initiation probability of mRNAs encoding mitochondrial proteins is somehow increased when those mRNA encounter a mitochondrion. We demonstrated that our model is consistent with the relationship between MLR and gene-specific estimates of ribosome occupancy and average number of ribosomes. We then further explored our model with a simple simulation of mRNA diffusion, entrapment and mitochondrial association which was able to reproduce trends seen in real data.

### Alternative Models

In this paper we report that translation initiation promoting sequence features correlate with a *low* degree of MLR — a surprising finding given that most mechanisms proposed for anchoring mRNA to the mitochondria require translation. As summarized above, we proceeded to incorporate that observation into a new mechanistic model of mRNA mitochondrial localization and explore the consistency and implications of that new model. Our simulation assumes that the mobility of translating mRNA is lower than that of non-translating mRNA. However this plausible assumption has not been firmly established. Recently exciting new developments in single-molecular imaging (81, 82, 83, 84, 85, 86) have addressed this question in mammalian cells. Some of these found ribosome load reduces mRNA mobility significantly (81, 84), but others reported the effect to be marginal (82, 83, 86). Hopefully future experiments will help clarify which is the case for the system we study here, but at this point we must consider both possibilities.

One alternative (non-mutually exclusive) explanation for the correlation between MLR and translation initiation promoting sequence features is that it reflects an evolutionary adaption to fit the needs of genes encoding (preferentially) co-translationally translocated protein products. The mitochondrial import machinery has a substantial but finite capacity, e.g. recent structural studies indicate that the TOM protein complex can simultaneously import no more than three polypeptide chains (87). Thus one could imagine that co-translationally translocated proteins have evolved modest translation initiation speeds to avoid locally overloading the import machinery with too many nascent peptides at one time. This explanation borrows closely from the logic of Quenault et al. (88), who suggested that mitochondrially associated Puf3 may facilitate co-translational protein import in the same way by slowing translation. Importantly, this alternative explanation does not directly contradict our new model, and the two explanations may both apply to some degree.

Evolutionary history and/or evolutionary adaption has also been evoked to explain other correlations between various gene features and MLR. For example, high MLR value genes tend to be of prokaryotic origin (7); and are enriched for components of certain mitochondrial inner membrane protein complexes, possibly facilitating the timely assembly of those complexes (19, 89). One could imagine that it is beneficial for the components of such complexes to delay the start of translation until after arriving at the mitochondria where they are assembled, and as a consequence they evolved to have slow translation initiation context (weak Kozak sequences etc).

Finally, as outlined in the introduction (see also reviews (90, 91, 92)) much progress has also been made in elucidating factors, such as Puf3, OM14, etc., involved in the mechanism of mRNA mitochondrial localization. Our results do not contradict or replace those observations, but rather offer another piece to the complex puzzle of understanding the mechanisms and significance behind mitochondrial mRNA localization.

## Supporting information

Supplementary Manuscript

## Acknowledgements

We thank Noriyuki Sakiyama, Kentaro Tomii and Toutai Mitsuyama for helpful comments during the early stages of this project.

