## Supplementary Manuscript for "Hallmarks of slow translation initiation revealed in mitochondrially localizing mRNA sequences"

### Supplementary Text for: Hallmarks of slow translation initiation revealed in mitochondrially localizing mRNA sequences

#### S1 Statistical Regression from mRNA features to MLR value

To investigate the relationship between mRNA features and MLR value we employed both linear regression and neural network models. To avoid overfitting, we partitioned each of the four microarray based datasets into equal sized training and test sets, and measured the correlation and mean average error (MAE) between predicted and actual MLR values on the test sets after training parameters on the training sets. For input features we tried each feature from table 1 in the main text, both as is and after taking the logarithm. In general, we found that the regression models produced equivalent or better results than the neural networks, and as the neural network results appeared subject to overfitting and varied with respect to network configuration and training, we used the regression models to evaluate the table 1 parameters. The neural network models served as a supplementary control to possibly catch non-linear relationships that linear regression (using standard and logarithmic parameter values) could not.

In agreement with Sylvestre et al. (1), we found a strong correlation between the logarithm of the ORF length and MLR values across all data sets (figure S1). These results also concur with a recent study by Williams et al. (2) which found that ribosome-nascent chains complexes of shorter protein products are significantly less likely to be observed at a mitochondrion. Therefore, we utilized the ORF length as a baseline for comparison with other predictors and calculated the correlation and mean average error (MAE) between predicted and actual MLR values for each of the four test sets (as shown in figure S1). Based on a comparison using MAE and correlation scores, we found that the ORF length performed better than any of the other predictors across all four datasets (see table S1 for a comparison with the best performing predictors). We also performed a multiple regression analysis using the ORF length and the table S1 variables to see if we could improve the localization prediction, and for each set we tested to see if there was a significant improvement (independent t-tests applied to the residuals from the baseline ORF length regression model and the residuals from each of the multiple regression models).

We note that a previous analysis by Weatheritt et al. (3) found a significant difference in the half-life between low and high MLR proteins. Although our results showed a low correlation between the protein half-life and MLR values, close inspection of the data revealed that mRNA with lower MLR values tended to have slightly longer half-lives than mRNA with high MLR value. The difference was non-linear, showing a small but sudden decrease in half-life near the mean MLR value. These findings concur with the Weatheritt et al. results, showing a modest but statistically significant non-linear effect, reflected in the larger magnitude of the rank correlation $\rho = -0.13$ than linear correlation $r = -0.05$ for protein half-life with MLR values in Table 1 of the main text.

#### S2 Search for Features which Can Complement Length in Predicting MLR value

Initially we explored many combinations of features, but were unable to find combinations of non-length features which consistently (across datasets) exceeded the prediction ability of log(ORF length) alone (results not shown). Thus we decided to search for features which could complement length in predicting MLR value. Although we tried combining each feature listed in Table 1, the only feature consistently able to complement length was the prediction score by TargetP (4) (shown) or MitoFates (5) (yielding similar results), which predict if an mRNA protein product possesses an N-terminal mitochondrial targeting signal. When combined with length, this MTS feature improved the regression performance over length alone in all of the microarray datasets; significantly (via independent t-tests applied to the residuals from the baseline length alone regression) in two of those, as shown in Supplementary Table S1. In other words, proteins imported into mitochondria via a non-MTS mechanism tend to have somewhat lower MLR values than MTS bearing proteins of similar length. We think this observation supports some role for a model in which co-translationally translocating nascent peptides can anchor their mRNA near the mitochondria. However the increase in regression performance gained from considering the MTS feature is not dramatic.

#### S3 Sampling of $5'$ UTR and $3'$ UTR Nucleotide Distributions

We computed the average MLR value of the set of mRNA with an `A`, `C`, `G`, or `T` at each position in the $5'$ UTR and $3'$ UTR respectively, as shown in the main text figure 4. Since a larger sample size typically results in smaller variance, it was important to ensure that similar sample sizes were used for each $5'$ UTR and $3'$ UTR position when comparing their variances. In particular, the $3'$ UTRs are on average longer and also tend to have more `A` nucleotides (reducing the `C`, `G`, and `T` counts). As a result, if we simply stipulate a large minimum sample size and compare the $5'$ UTR and $3'$ UTR, then the resulting samples would include many $3'$ UTR `A`s, but relatively few samples for other (nucleotide, position) pairs. On the other hand a small minimum sample size would result in throwing away a considerable amount of data. We therefore set a sample size range instead, using a minimum of 75 and maximum of 100 counts for each nucleotide position, where MLR values from any additional mRNA were not included when the sample size reached the maximum (in this case the first 100 samples were chosen from a randomly ordered dataset). We chose this sample size range as it allowed us to utilize most or all of the samples in each `A`, `C`, `G`, or `T` subset taken from 524 mRNA analyzed in total (e.g. 20% of the 524 mRNA might for example have a `C` at the first position of the $5'$ UTR, in which case 100 out of 105 samples are used). In addition, this also allowed us to maintain more than half of the data points in the main text figure 4, and the sampling became normalized such that the `A` and `T` subsets had the same sample size at all positions, while the `C` and `G` counts were almost exclusively higher in the $5'$ UTR (thus any potential remaining sample size bias should tend to increase the $3'$ UTR variance, but in fact the $5'$ UTR variance was significantly higher).

#### S4 Simulation of mRNA Diffusion to Mitochondria

We developed a simulation to model mRNA diffusion/entrapment and mitochondrial import to explore the possibility that changes in mRNA mobility upon translation initiation may explain some trends in mRNA mitochondrial localization by comparing simulated MLR values with the ORF length and with real MLR values. Our simulation models diffusion using a random walk in three dimensions as shown in figure S2, taking a set of gene translation

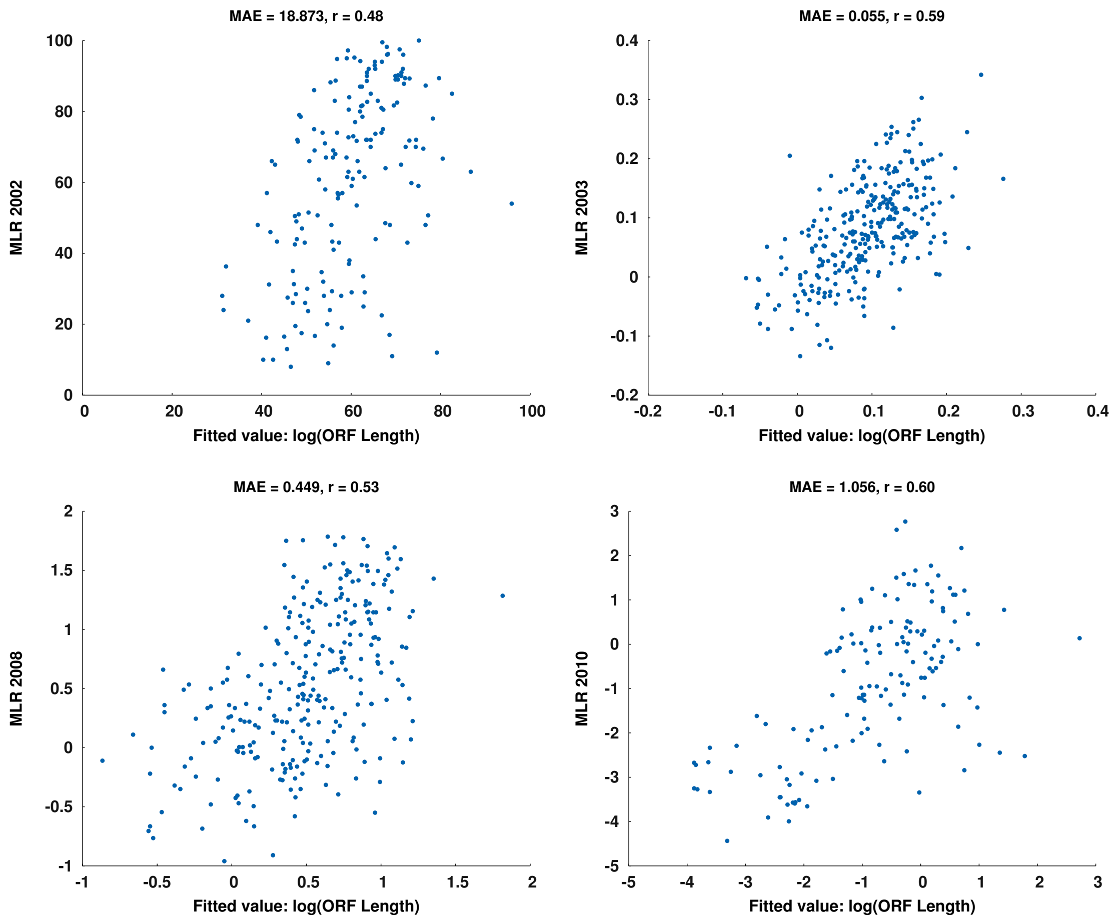


**Figure S1:** Plots for the ORF length regression model and MLR test datasets. The ORF length displayed the strongest correlation values and lowest mean average errors (denoted $r$ and MAE respectively) across all MLR datasets compared with other mRNA features.

| | **MLR2002** | | | **MLR2003** | | | **MLR2008** | | | **MLR2010** | | |
|---|---|---|---|---|---|---|---|---|---|---|---|---|
| | $r$ | MAE | $p$ | $r$ | MAE | $p$ | $r$ | MAE | $p$ | $r$ | MAE | $p$ |
| **ΔSsa1 affect** | .51 | 19.1 | .087 | .40 | .061 | .019 | .48 | .502 | .041 | .57 | 1.15 | .285 |
| **ORF Length** | .57 | 17.2 | — | .62 | .052 | — | .60 | .437 | — | .64 | 1.09 | — |
| **ΔSsa1 affect+ORF Length** | .62 | 16.2 | .082 | .63 | .051 | .104 | .63 | .450 | .771 | .71 | 1.00 | **.044** |
| **MTS Score** | .37 | 20.6 | .021 | .20 | .065 | .000 | .35 | .498 | .010 | .31 | 1.26 | .008 |
| **ORF Length** | .48 | 18.9 | — | .59 | .055 | — | .53 | .449 | — | .60 | 1.06 | — |
| **MTS Score+ORF Length** | .57 | 18.0 | **.030** | .61 | .054 | .118 | .57 | .433 | **.000** | .63 | 1.04 | .257 |

**Table S1:** Comparison between the logarithm of the ORF length and other features that were investigated for MLR prediction. MTS denotes the N-terminal mitochondrial matrix signal prediction score of the protein product. $p$ denotes the statistical significance of differences in MLR prediction performance using the features indicated for each row, compared to using ORF length alone.

initiation probabilities as input and producing simulated MLR values as output. The simulation code can be found at: `https://github.com/gitmitosim/mitosim`. We also performed a companion follow-up study producing gene specific simulated MLR values, as described in a separate document.

#### S5 Summary of Discretionary Simulation Parameters

We simulated mRNA for the set of genes from Sylvestre et al. (1), the largest of the four primary MLR datasets. To simulate an mRNA diffusion process within a yeast cell, we mod-

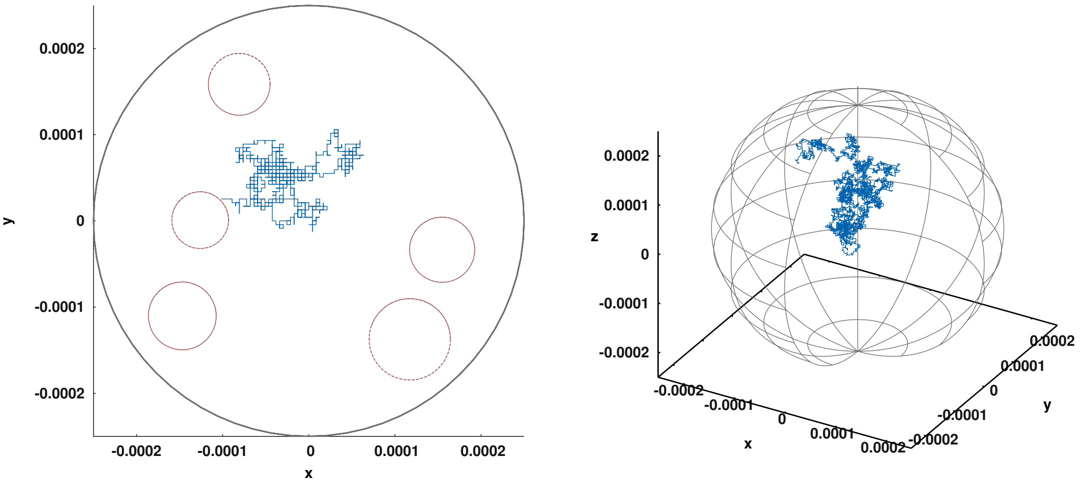


(a) (b)

**Figure S2:** The simulation was performed in three dimensions but the above examples shows both an example of a 2D (left) and 3D (right) simulation for illustration purposes. The blue line shows the course traveled by an mRNA molecule (starting at the center of the cell), and red circles denote mitochondria (not shown in 3D example for sake of visual clarity of the mRNA course).

| **Quantity** | **Value** | **Source** |
|---|---|---|
| Radius of cell (sphere) | 2.5$\mu$m | Study of yeast size and growth rate (9) |
| Number of mitochondria | 5 | Measurement in yeast (10) |
| Radius of mitochondria (sphere) | 0.25–0.5$\mu$m (sphere) | Measurements under different conditions (11) |
| mRNA Diffusion coefficient (non-translating) | $1.23 \times 10^{-9}$ cm$^2$/s | Consideration of published estimates (6–8) |
| mRNA Diffusion coefficient (translating) | $0.30 \times 10^{-9}$ cm$^2$/s | Consideration of published estimates (6–8) |

**Table S2:** Values of parameters used in the simulation.

eled both the cell and mitochondria as spheres and assumed that all mRNA start diffusing at the center of the cell at the beginning of the simulation. The simulation was modeled using the physiological parameters described in Table S2. The diffusion coefficients used in our simulation are in the same ball park as published estimates (6–8) which vary considerably.

The remaining parameters needed for our simulation are: the size and number of time steps to perform per random walk, the number of random walks to perform for each gene, the translation initiation probability of each gene, the probability of loss of mobility via cytoskeletal entrapment of mRNA molecules before and after initiating translation, the multiplicative boost in translation initiation probability experienced when approaching a mitochondria, and the probability that a translating mRNA will anchor when contacting a mitochondrion.

We adopted 0.01 seconds as the span of each time step in the simulated random walks, which is short enough that at each step an mRNA molecule will move by only a small fraction of the radius of the cell (the simulation performs a random-walk such that each molecule randomly moves along the x, y, or z axis at every time step. We used a total simulation time of 660 seconds to match the median half-life of yeast mRNA reported by Miller et al. (12). Given the stochastic nature of a random walk, to obtain stable results multiple runs must be averaged for each gene. Here, we set the total number of mRNA to 1000 as this appeared to be a good balance between reasonable simulation computation time and reducing the variance between estimated MLR values across repeated simulations.

We derived gene-specific translation probabilities based on average time to translation initiation times reported by Shah et al. (13). We reasoned that the average time to initiation translation would naturally be modeled by an exponential distribution over continuous time, which can be well approximated by a discrete geometric distribution over time steps if the span of each step is relatively small. Following a geometric distribution,

$$\bar{N}_I^g = \frac{1}{P_I^g}, \quad P_I^g = \frac{1}{\bar{N}_I^g} \tag{1}$$

where $\bar{N}_I^g$ is the average number of 0.01s time steps before translation initiation according to the Shah et al. data and $P_I^g$ denotes the gene-specific probability of translation initiation during the span of a single time step.

As described in the main text, our model assumes that there is a mechanism that promotes translation in the vicinity of mitochondria, that an mRNA may become entrapped in the cytoskeleton, and that the probability of entrapment is higher if an mRNA has initiated translation. In practice, the entrapment probability would be gene-specific, but for simplicity we employ a single probability of entrapment at each time step (denoted by $p_{E_I}$) for all mRNA molecules that have initialized translation, and another (smaller) entrapment probability ($p_{E_U}$) for uninitiated mRNA molecules which have not yet initiated translation. We use $x$ to denote the ratio of these two entrapment probabilities.

The entrapment probability $p_{E_U}$ is derived for a given value of $x$ by performing a grid search with a smaller number of mRNA in the simulation (the search uses 100 for each gene instead of 1000 as in the 'full-scale' simulation). The grid search starts with an estimated value for $p_{E_U}$, and then iteratively alters $p_{E_U}$ until approx. 40% of mRNA become entrapped as reported by Fusco et al. (6) (where the percent of entrapped mRNA is calculated as an average across all genes we simulate for). The $p_{E_U}$ value found is then used in the full simulation, where experiments with values of $x = 5, 10, 15, 20$ gave qualitatively very similar results (not shown).

Finally, we note that in our simulation mRNA always initiate translation upon reaching a mitochondrion (translation is not re-initiated if started prior to localization) and there is then a probability of anchoring given by $p_A$. For the simulation in the main text we set $p_A = 1.00$ but as can be seen in table 3, using different values of $p_A$ showed little effect on results (see discussion in main text).

In our resulting simulation, mRNA localization is thus dependent on when an mRNA initiates translation such that translating mRNA have slower diffusion rates (see table S2) and a higher chance of entrapment as described above. The simulation thus investigates if these two mechanisms could account for the correlation between translation initiation based on the assumptions we have described here.

The parameters discussed in this section are summarized in table S3.

| **Param.** | **Description** | **Value** |
|---|---|---|
| $N_{\text{mRNA}}$ | Number of mRNA random walks simulated per gene | 1000 |
| $\delta_T$ | Length of each simulation time step | 0.01 seconds |
| $T$ | Total time of simulation | 660 seconds |
| $N_S$ | Number of simulation steps | 66000 |
| $p_{E_U}$ | Base entrapment probability | $9 \times 10^{-5}$, ($7 \times 10^{-6}$–$9 \times 10^{-5}$) |
| $p_{E_I}$ | Entrapment probability for translating mRNA | $p_{E_U} \cdot x$ |
| $x$ | Fold increase of above probability upon translation | 5, 1–20 |
| $p_A$ | Probability of anchoring to Mitochondria | 1.0, (0.05–1.0) |

**Table S3:** Summary of discretionary simulation parameters and their values.

#### S5.1 Simulation Runs

We ran the simulation with values $x = 5, 10, 15, 20$, and for each $x$ value the simulation consisted of the steps shown as shown below:

1) Use the average initiation probability (and time) from Shah et al. (13) for all genes, set $x$ to whichever value we are using in the current simulation, and stop diffusion of an mRNA if it reaches a mitochondrion.

2) Perform grid search using 100 mRNA for each gene to find $p_{E_U}$

3) For each gene: run simulation again with derived $p_{E_U}$ value and use the initiation probability for mRNAs associated with each gene.

Simulations were run ten times to compare variability between results, and the $x$ value that best fit with the Fusco et al. (6) observations was selected. Simulation output was consistent and the average correlation between simulated and real MLRs across simulations for $x = 5$ was $r = 0.44$. All simulations were qualitatively similar with s.d. $= 0.021$ where Figure 7 in the main text shows the one with the correlation closest to the average.

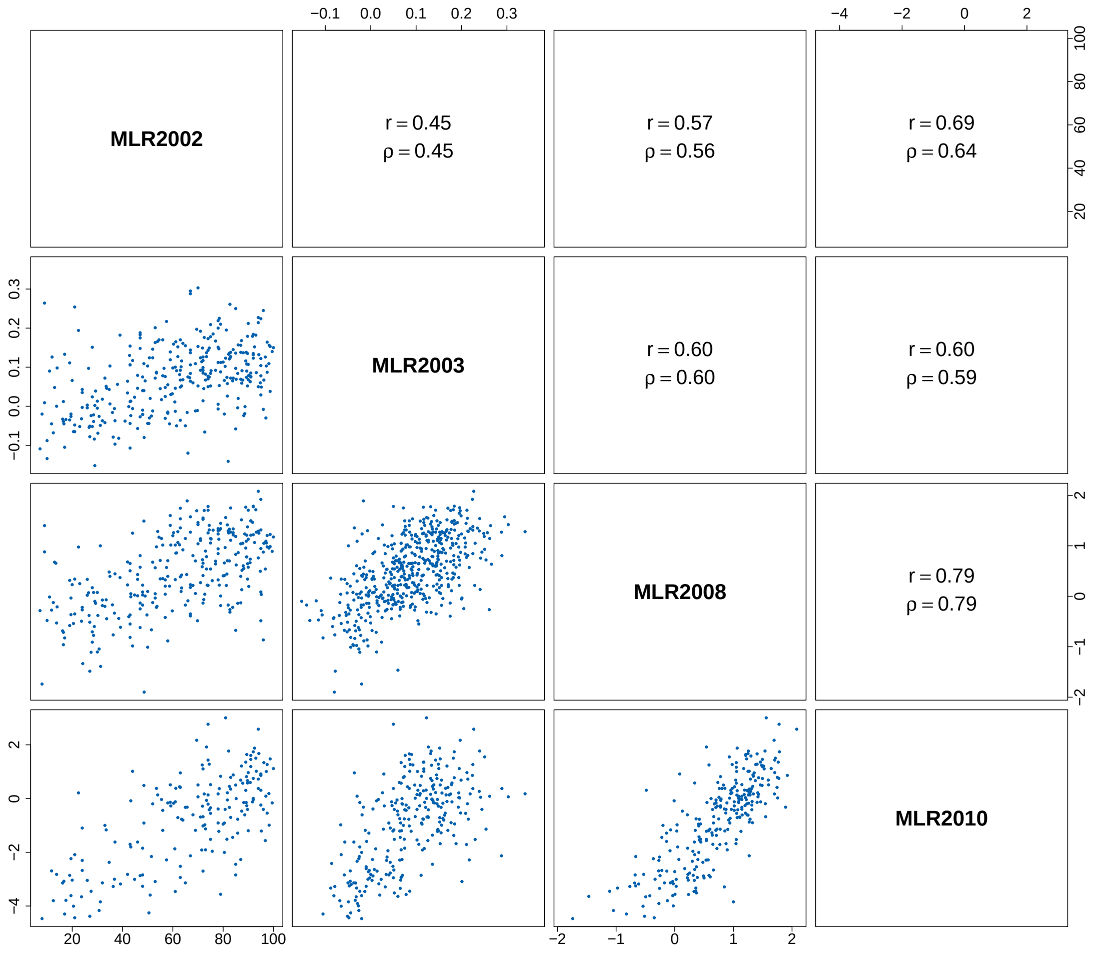


**Figure S3:** Scatter plots of MLR dataset pairs with Pearson and Spearman correlation values shown as $r$ and $\rho$ respectively. Although the four different MLR sets that we analyzed utilized different scales, the linear correlation between them was generally strong.

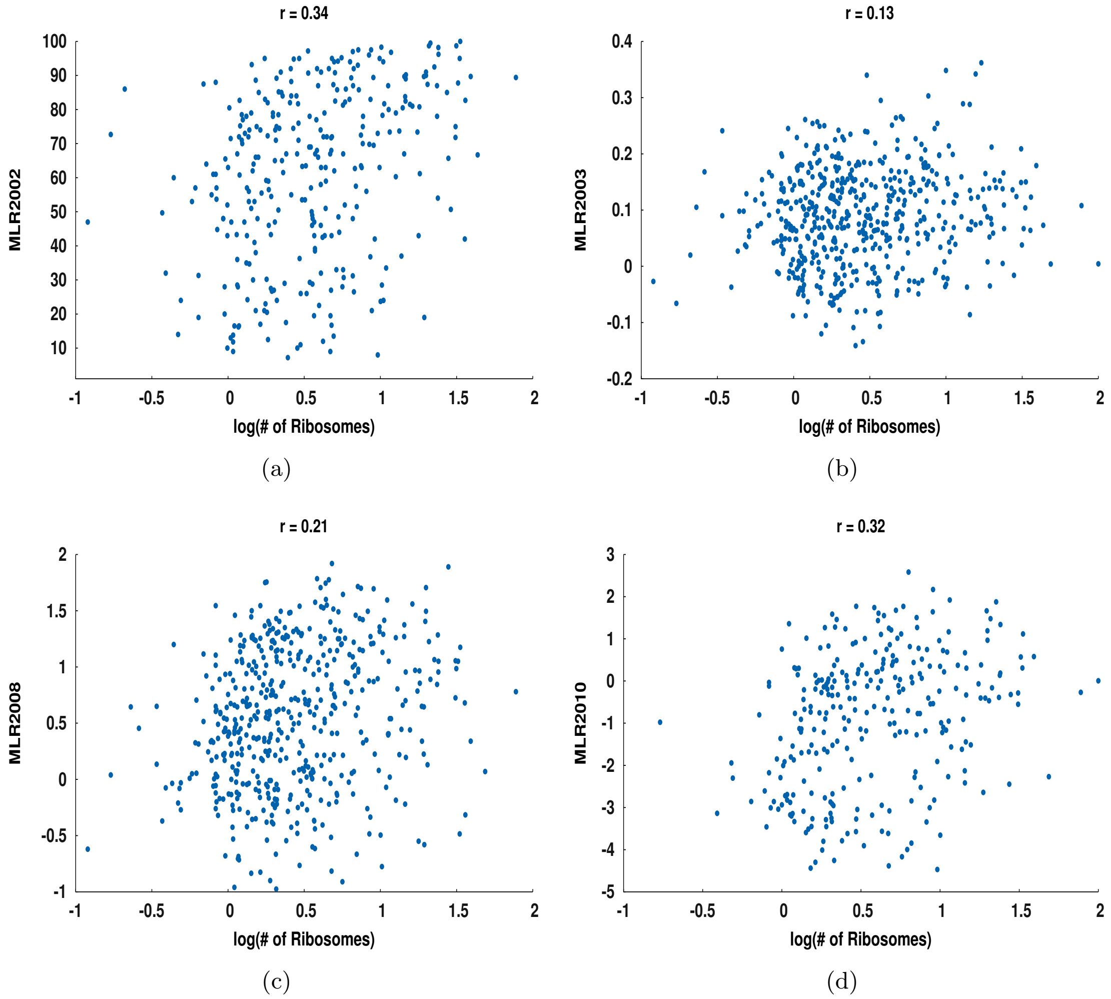


**Figure S4:** Relationship between the log(# of ribosomes) and the four MLR datasets: a) MLR2002, b) MLR2003, c) MLR2008, d) MLR2010.

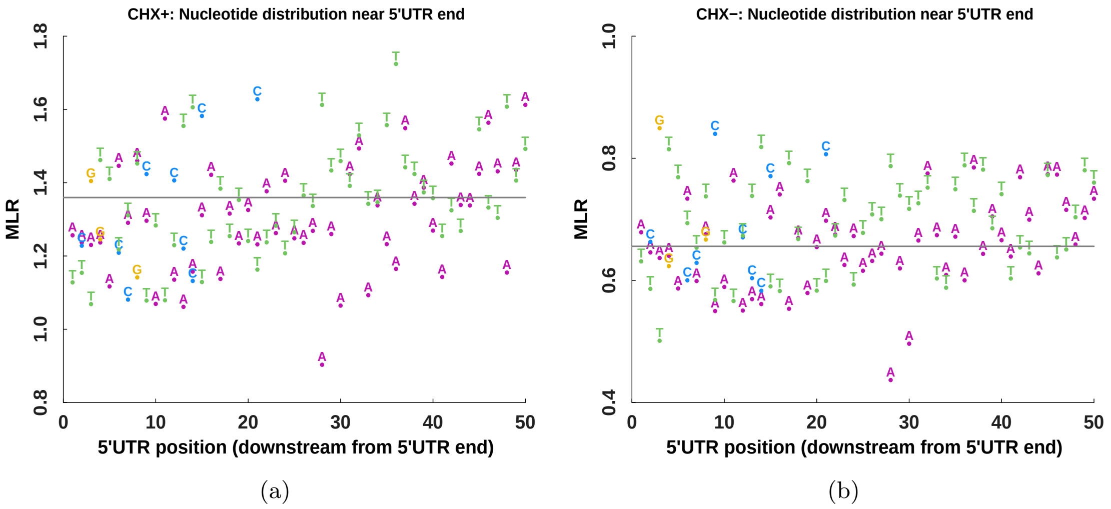


**Figure S5:** Average MLR values of individual nucleotides in the -10 to -1 region upstream of the start AUG codon are shown for the CHX+ and CHX- datasets.

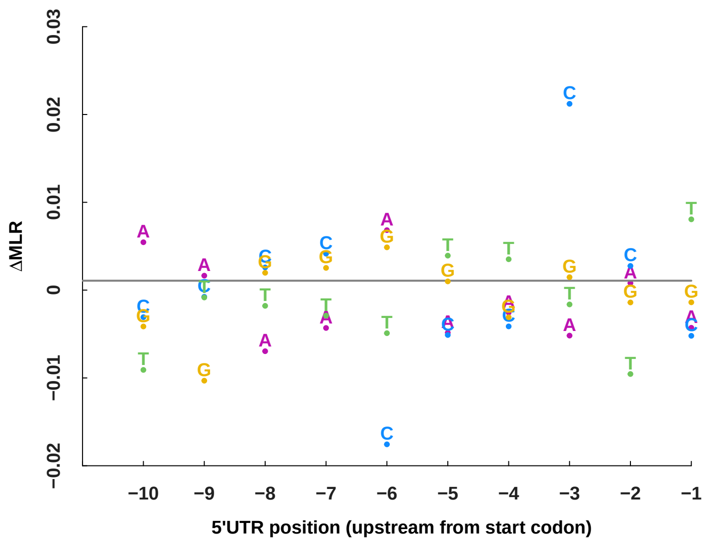


**Figure S6:** Difference between the MLR4 values of individual nucleotides in the -10 to -1 AUG region of mRNA with and without an upstream AUG.

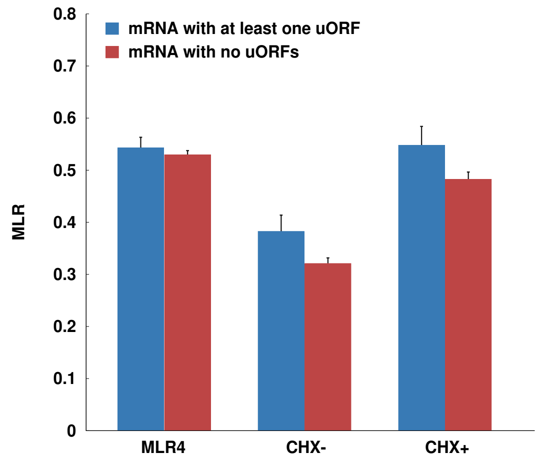


**Figure S7:** The average MLR values of genes with mRNA containing at least one uORF (blue bar) or none (red bar) for the MLR4, CHX+, CHX- datasets. Error bars show standard error of mean estimation.

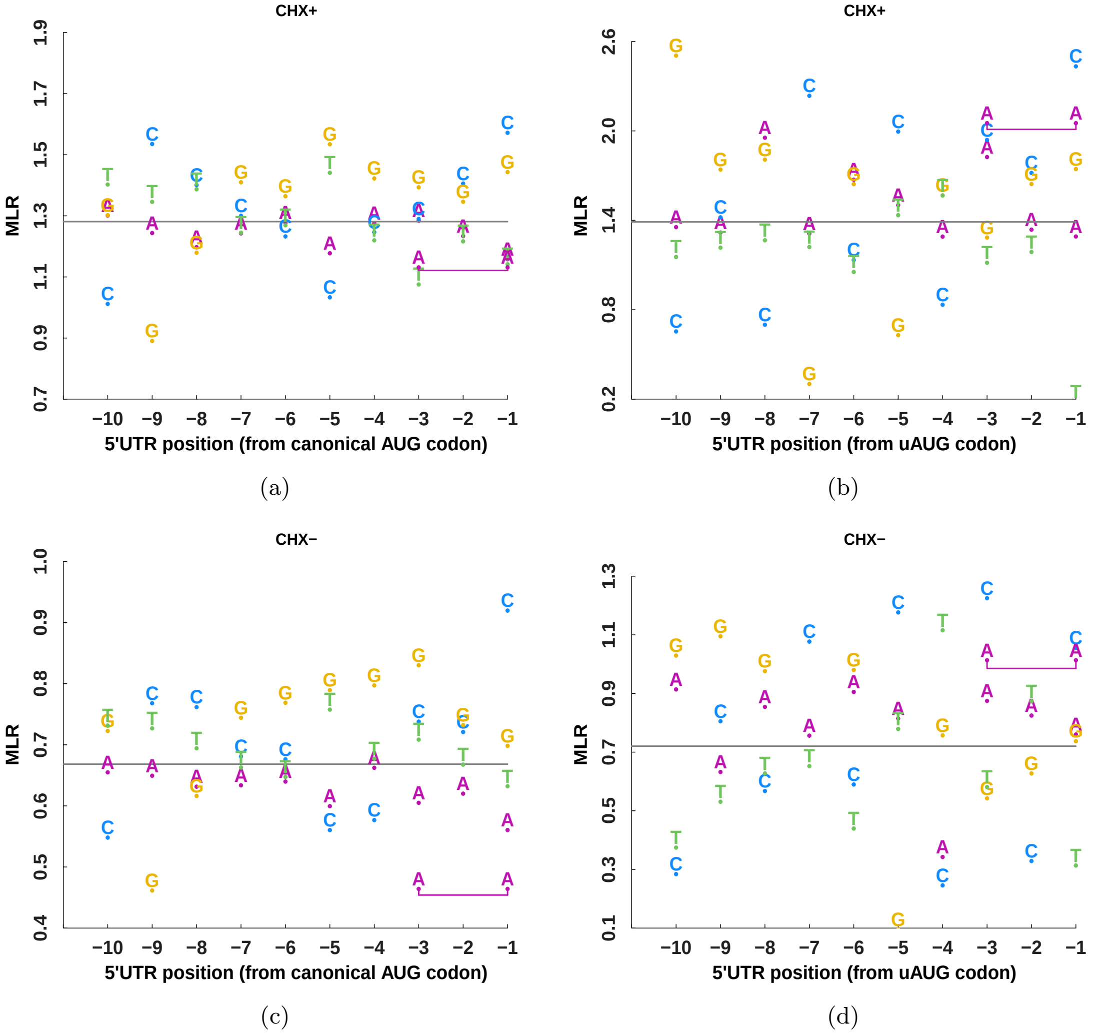


**Figure S8:** Comparison of context correlation with MLR value for canonical `AUG`s (left half) versus the `AUG`s of single uORF mRNAs (right half). The two horizontal rows show results for the CHX+ and CHX- datasets respectively. For each case, the average MLR value for mRNA with an `A`, `C`, `G`, or `T` respectively, at the -10 to -1 positions upstream of the `AUG` are shown. The bar linking the -1 and -3 `A`s represents the average MLR for mRNAs with `A`s in both those positions.

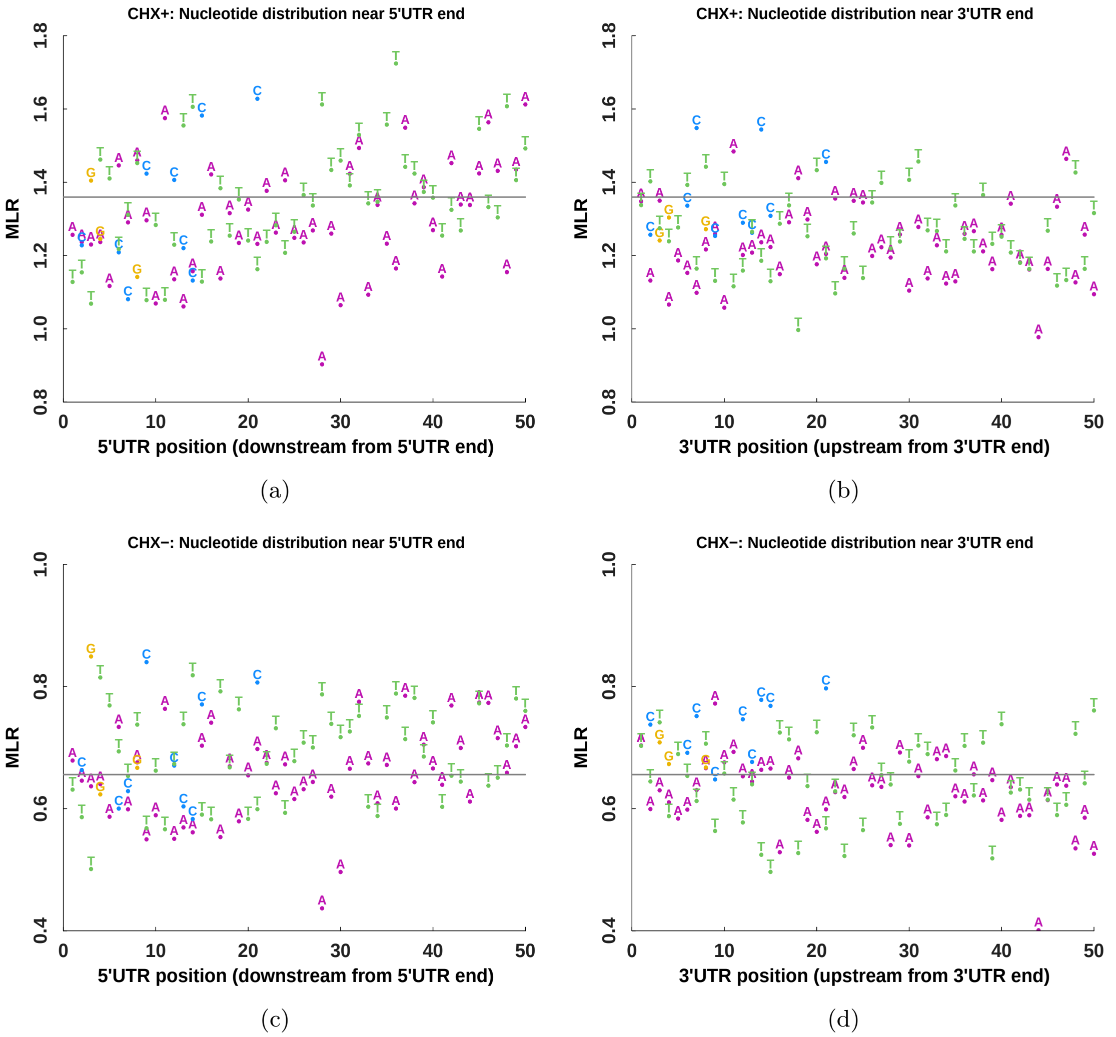


**Figure S9:** The variance in average MLR value conditioned on (nucleotide, position) pairs is greater for the 5'UTR than the 3'UTR end (only nucleotides with sufficient counts shown, see text for details)

.

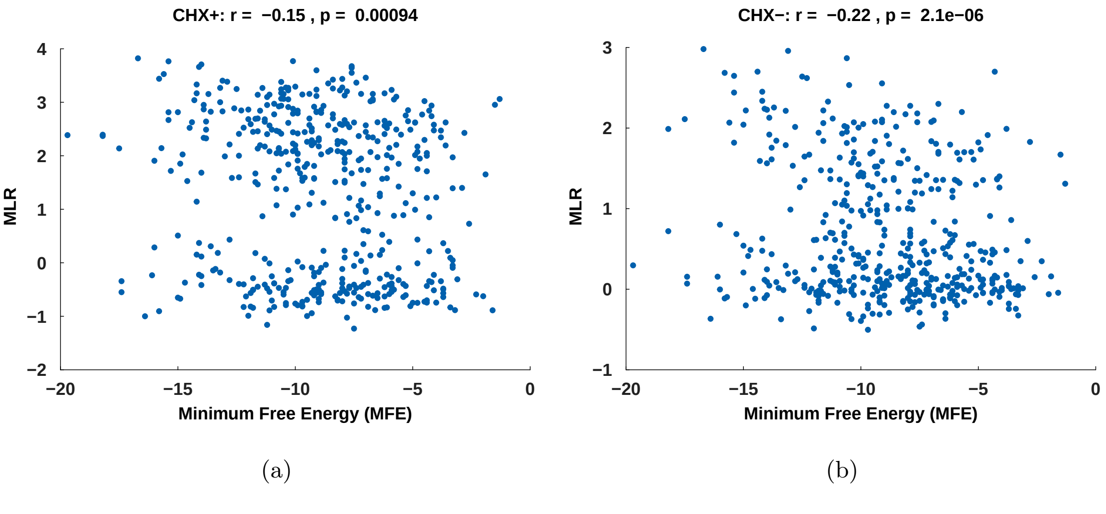


**Figure S10:** Scatter plots of the calculated minimum free energy of the -15 to +50 region of mRNA and their MLR values from the CHX+ and CHX- datasets are shown. $r$ denotes the Pearson correlation coefficient and $p$ its statistical significant by $t$-test against a null hypothesis of no correlation.

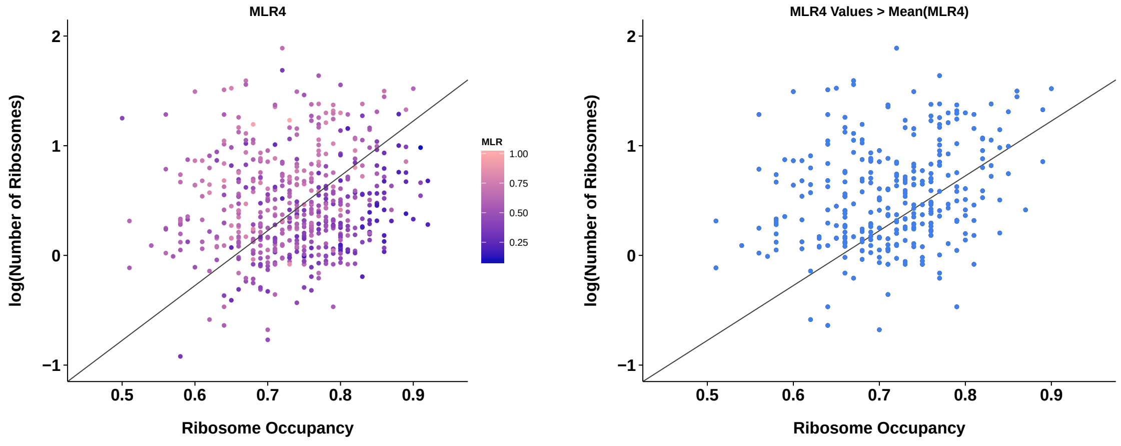


(a) (b) (c)

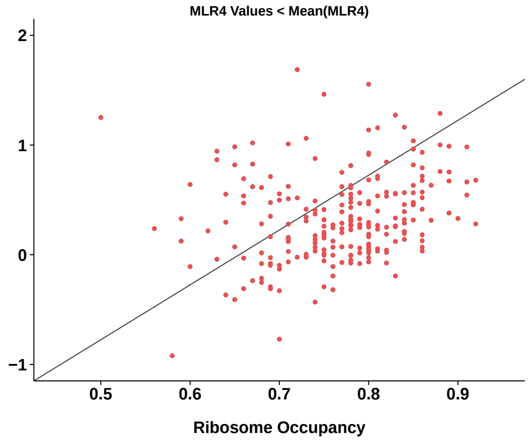


(d)

**Figure S11:** Relationship between ribosome occupancy and the number of ribosomes (points color graded by MLR4 values). Line added for visualization purposes to show relative position of different points in each figure: a) All points in dataset. To illustrate separation between points based on MLR values: b) Points with MLR values above the mean MLR value (= 0.53). c) Points below the mean MLR.

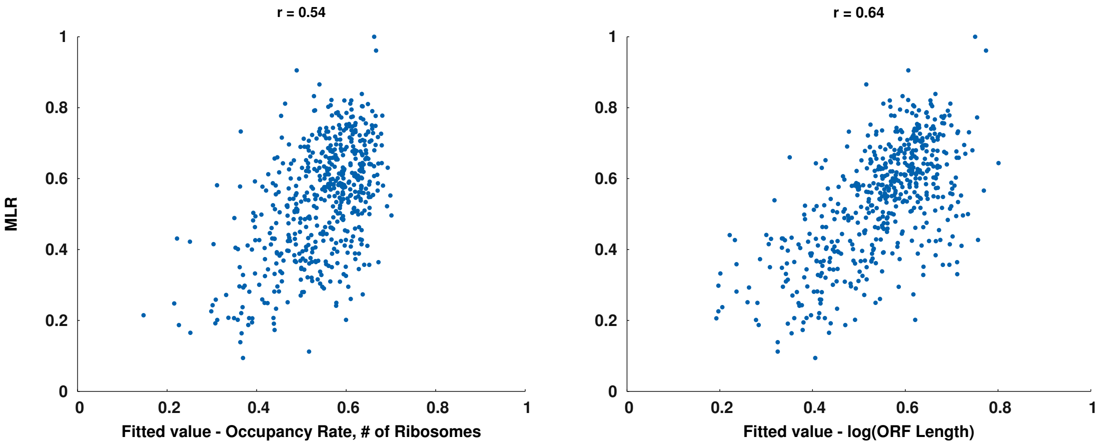


(a) (b)

**Figure S12:** a) Neural network model using the occupancy rate and ribosomal counts to predict MLR4 values. b) Neural network model that uses the ORF length to predict MLR4 values.

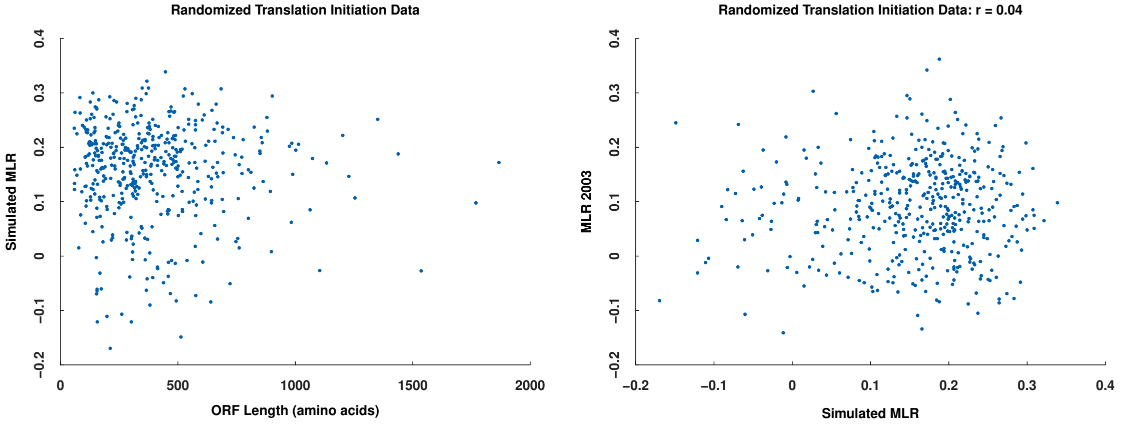


(a) (b)

**Figure S13:** For each gene we computed a simulated MLR value using randomly generated translation initiation times. a) The relationship between simulated MLR values and the ORF length was not similar to the real data shown in Figure 7c). b) The simulated MLR values showed a poor correlation with real MLR data.
